# Nerve-targeting and pain-promoting transcriptomic signatures in early Guillain-Barré syndrome

**DOI:** 10.1101/2025.09.24.678413

**Authors:** Jayden A. O’Brien, Joseph B. Lesnak, Ishwarya Sankaranarayanan, Asta Arendt-Tranholm, Nikhil Inturi, Spoorthi Sadasivuni, Hemanth Mydugolam, Katelyn E. Sadler, Theodore J. Price, Eroboghene E. Ubogu

**Author notes:** Co-corresponding authors. T.J.P.: 800 W Campbell Rd, BSB 14.102, Richardson, TX 75080, USA. E.E.U.: 1720 7th Avenue South, Sparks Center Suite 200 Birmingham, AL 35244, USA. **Conflict-of-interest statement:** The authors declare no competing interests.

## Abstract

Guillain-Barré syndrome (GBS) is an autoimmune disorder that causes weakness, sensory loss, autonomic dysfunction, and chronic neuropathic pain. The mediators responsible for driving early autoimmune injury in the most common GBS variant, acute inflammatory demyelinating polyradiculoneuropathy (AIDP), remain incompletely understood. We performed single-cell and bulk RNA sequencing on peripheral blood mononuclear cells collected from early untreated AIDP-variant GBS patients and healthy controls to comprehensively deduce leukocyte transcriptome alterations and predict disease- and pain-driving interactions between pathogenic leukocytes and peripheral nervous system cells. We found that classical, intermediate, and non-classical monocytes were expanded and upregulated genes associated with type I and II interferons, JAK/STAT signaling, and NLRP3 inflammasome engagement. CD8+ T cells were highly proliferative and likewise upregulated JAK/STAT signaling. *CD4+FOXP3+* regulatory T cells upregulated *PRDM1* and *CD74* in a signature that may indicate functional exhaustion. A subpopulation of highly activated intermediate monocytes upregulated genes related to angiogenesis and oncostatin M. Differential expression-based cell-cell interaction analysis between GBS leukocytes, Schwann cells, and sensory neurons predicted engagement of ligand-receptor pairs with nerve integrity and pain functions, including epiregulin, interferon-beta, adrenomedullin, clusterin, IL-6, IL-15, and CCL4. Functional validation demonstrated that CCL4 sensitizes human sensory neurons *in vitro*. These results unearth molecular interactions by which specific leukocyte populations in AIDP-variant GBS may participate in peripheral nerve injury and drive neuropathic pain. Many of these targets may be amenable to therapeutic modulation using available approved and investigational drugs, potentially providing drug repurposing opportunities.

## Introduction

Guillain-Barré syndrome (GBS) is a peripheral nerve-restricted autoimmune disorder causing muscle weakness, sensory loss, and autonomic dysfunction (1). GBS results in death in 3-5% of patients, and survivors often experience significant morbidity in the form of chronic neuropathic pain, residual sensory and motor deficits, loss of prior functional capabilities, and employment loss (2). The most common variant of GBS worldwide is acute inflammatory demyelinating polyradiculoneuropathy (AIDP), which, unlike acute motor axonal neuropathy, is primarily driven by injury to myelin sheaths surrounding peripheral axons, termed demyelination (3). Despite significant advances in the pathogenesis, diagnosis, and treatment of GBS, major knowledge gaps exist, such as the relative importance and interactions of different effector cells and their signaling pathways at different phases of this tissue-specific autoimmune disorder. Molecular mimicry is considered critical to the pathogenesis of GBS, involving the generation of pathogenic antibodies against neuronal gangliosides or peripheral nerve myelin proteins or lipids (4, 5). However, the pathogenic changes that are definitively responsible for AIDP-variant GBS remain elusive.

Based on cytokine profiling, *in vitro* T cell screening, single-cell RNA sequencing and T cell receptor (TCR) sequencing, there is emerging evidence that AIDP-GBS is associated with autoreactive effector memory CD4+ T cells with a T-helper 1 (Th1)-like phenotype that target peripheral nerve myelin antigens, implying an essential pathogenic role in this autoimmune disorder (6–9). Histopathological evaluation of biopsies from AIDP-variant GBS patients indicates hematogenous leukocyte trafficking, with monocyte/macrophage, CD4+ and CD8+ T cell, and B cell/plasma cell infiltration associated with demyelination and axonal degeneration (10–15).

Monocytes/macrophages are the most common leukocyte subpopulation in AIDP peripheral nerves and in an experimental autoimmune neuritis (EAN) model, where they are observed to invade the myelin sheaths of structurally intact axons (16). This is hypothesized to be a response to specific antibody-antigen complexes on the myelin surface, on Schwann cells – the supportive glial cells of the peripheral nervous system, a subset of which are responsible for myelination – or at the node of Ranvier between myelin segments (3, 17–19). Activated monocytes and tissue-resident macrophages may further enhance the pathogenic immune response, worsen local inflammation, and induce axonal injury via cytokine and other pro-inflammatory molecule release.

T cells are the next most prevalent leukocyte subpopulation in peripheral nerve biopsies in AIDP and EAN, with CD4+ T-helper cells (polarized to Th1 and Th17 phenotypes) implicated in pathogenesis via the production of pro-inflammatory cytokines and chemokines in peripheral nerves and nerve roots that enhance inflammation and activate macrophages. T cells increase their expression of interferon-γ (IFNγ) in response to stimulation with ganglioside GM1 in vitro (20). More recently, single-cell RNA sequencing (scRNA-seq) and T cell receptor (TCR) sequencing of circulating T cells in GBS has demonstrated the existence of myelin protein-specific CD4+ Th1 and CD8+ T cells in blood, cerebrospinal fluid, and peripheral nerve (9). In addition, decreases in the number of circulating CD4+CD25+FoxP3+ regulatory T cells have been shown during the acute phase of GBS (3, 21, 22).

We aimed to comprehensively characterize early untreated AIDP-variant GBS patient peripheral blood mononuclear cells (PBMCs) by taking advantage of major technological and bioinformatics advances in transcriptomics and predicting interactions with nerves that may drive disease and inform new therapeutic targets. We identified key early-stage pathogenic effector leukocyte subpopulations and discovered signaling pathways, networks, and interactions with Schwann cells and sensory neurons that may be suitable for therapeutic modulation using a small but well-characterized AIDP-variant GBS patient cohort. We then aimed to determine which signaling molecules upregulated in specific AIDP-variant GBS leukocyte subsets are predicted to induce peripheral nervous system pathology by intersecting the GBS data with existing transcriptomic atlases of human peripheral nervous system tissues (23–27). Finally, we validated one of these potentially pronociceptive interactions between GBS patient leukocytes and peripheral sensory neurons and Schwann cells using functional assays.

## Results

GBS and control participant ages were adequately matched (GBS: mean ± SD 39.3 ± 14.8; control: 38.3 ± 12.4). Two out of four GBS patients had known inflammatory exposures (pancreatitis based on elevated pancreatic enzymes with no detected cause; and infectious gastroenteritis of unknown pathogenic origin); the remaining two had no reported infection or infectious contacts prior to disease onset. Key demographic and clinical information of included participants is presented in **Table 1**. Complete clinical characterization is available in **Supplemental Table 1**.

**Table 1.** Participant demographic and clinical information. Patients with a designation of early AIDP-variant GBS (*n* = 4) and age- and sex-matched healthy controls (*n* = 3) were included. CSF glucose was normal (40-75mg/dL) in all GBS patients. ^Outside normal range (complete blood count: 4.0-11.0×10^3^/cm^3^; CSF protein: 18-52mg/dL; CSF leukocytes: 0-5 cells/cm^3^).

| Subject Code | Classification | Age | Sex | Race | Symptom Onset (days before blood draw) | Reported Exposures (time prior to symptom onset) | Prior Medical History | Electrodiagnostics | Complete Blood Count ($10^3/\text{cm}^3$) | CSF Protein (mg/dL) | CSF Leukocytes (count/$\text{cm}^3$) |
| --- | --- | --- | --- | --- | --- | --- | --- | --- | --- | --- | --- |
| GBS1 | Early GBS | 29 | M | White | 6 | High work-related stress (3 weeks) | None reported | Mild-moderate acute polyradiculopathy | Normal | 79<sup>^</sup> | 0 |
| GBS2 | Early GBS | 42 | F | Black | 7-14 | Pancreatitis (4-6 weeks) | Chronic gastritis | Mild-to-moderately severe acute polyradiculopathy with absent median and ulnar F-wave responses | Normal | 31.2 | 3 |
| GBS3 | Early GBS | 27 | F | White | 12 | None | Cerebral atrophy with developmental delay, diffuse spondylosis | Moderately severe acute, non-length-dependent, distally predominant, primarily demyelinating sensorimotor polyradiculoneuropathy with secondary axonal involvement | Not reported | 71<sup>^</sup> | 0 |
| GBS4 | Early GBS | 59 | M | White | 6 | Gastroenteritis (7-10 days) | Hypertension | Moderate-severe acute primarily demyelinating sensorimotor polyradiculoneuropathy with secondary axonal involvement | 21.94<sup>^</sup> | 128<sup>^</sup> | 13<sup>^</sup> |
| HC1 | Control | 46 | M | Black | - | None | - | - | - | - | - |
| HC2 | Control | 24 | F | White | - | None | - | - | - | - | - |
| HC3 | Control | 45 | M | White | - | None | - | - | - | - | - |

### Bulk transcriptomics shows myeloid and CD8+ cell responses in untreated GBS patient PBMCs

To characterize the overall changes present in key leukocyte populations in early AIDP-variant GBS patients compared to age- and sex-matched controls, bulk RNA sequencing was conducted on PBMC samples sorted for myeloid cells (CD11b+), cytotoxic T cells (CD3+CD8+), and helper T cells (CD3+CD4+) and underwent differential expression analysis. The complete bulk RNA sequencing differential expression results are presented in **Supplemental Table 2**.

There were 164 upregulated and 44 downregulated genes in CD11b+ cells (**Figure 1A**). These genes were most significantly involved in type I interferon-mediated antiviral signaling (*IFNB1, IFIH1*, *IFIT1, IFIT2, IFIT3, IFITM3, GBP2, OAS1, OAS2*), IFNγ signaling (*STAT1*, *SOCS3*), and related cytokines and chemokines (*CXCL9*, *CXCL10*, *IL6*, *IL15, IL31RA*). *FAS* and IL-27 signaling pathways were also upregulated, as were *ADM*, *MARCO*, *CCR5, CLU,* and *EREG* (the latter encoding the growth factor epiregulin; **Figure 1B-C**).

**Figure 1.**
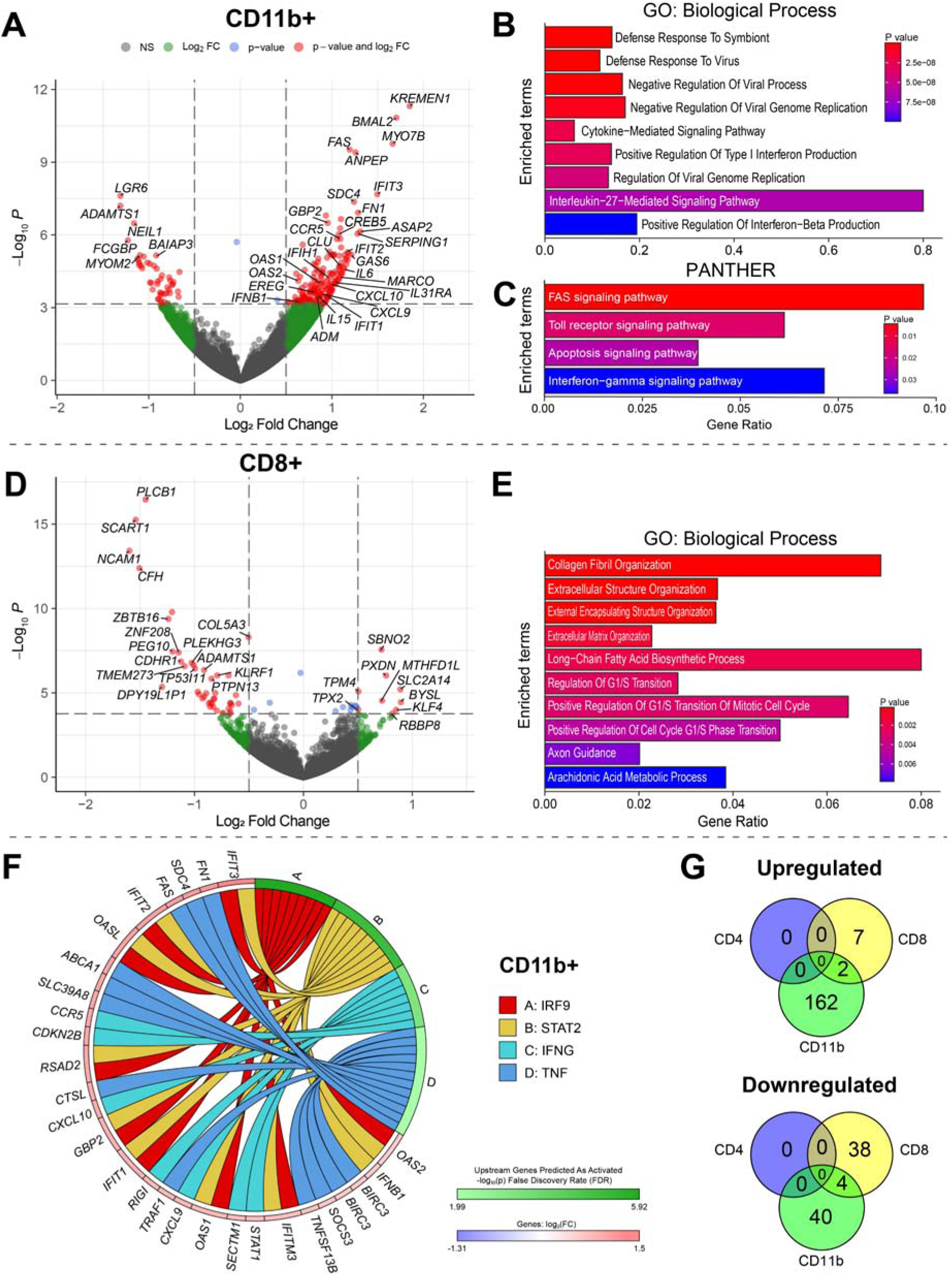
Bulk RNA sequencing of cell-sorted CD11b+, CD8+, and CD4+ cells shows transcriptomic changes related to type I and II interferon production, FAS signaling, and cell proliferation in early AIDP-variant GBS. **(A)** Volcano plot of differentially expressed genes in CD11b+ myeloid cells. Positive fold-changes indicate genes with higher expression in GBS patients than in healthy controls. Log_2_FC > |±0.5| and P_adj_ < 0.05 were used as cutoffs. **(B)** Gene set enrichment analysis using gene ontology (GO) annotations in Enrichr showed that the most overrepresented biological processes in myeloid cells included viral response, type I interferon signaling, and cytokine production. **(C)** Further gene set enrichment using PANTHER indicated FAS and interferon-γ as among the most highly overrepresented signaling pathways. **(D)** Volcano plot of differentially expressed genes in CD8+ sorted cells. **(E)** The most highly overrepresented biological processes included those related to mitosis and extracellular matrix organization. **(F)** Genes differentially expressed in the CD11b+ cell portion were significantly associated with the activity of upstream regulators *IRF9*, *STAT2*, *IFNG*, and *TNF* (all *P* < 0.05). Pathways were generated using iPathwayGuide. **(G)** The upregulated and downregulated genes in CD11b+, and CD8+ sorted cells were largely unique to each population. There were no differentially expressed genes in CD4+ sorted cells in the bulk RNA sequencing data.

The differentially expressed genes in CD8+ sorted T cells are presented in **Figure 1D**. The alterations in CD8+ T cell genes were most significantly associated with biological processes pertaining to the extracellular matrix (including type V collagen genes *COL5A1* and *COL5A3*), and mitosis (**Figure 1E**). This CD8+ T cell phenotype suggests a high level of expansion along with increased capabilities for intercellular interactions.

Given the large number of differentially expressed genes in CD11b+ cells in GBS, we analyzed the predicted upstream regulators of these genes. This provides information on the predicted presence of proteins that are consistent with the pattern of upregulated genes found in these cells. The top predicted regulators of CD11b+ cells were IRF9, STAT2, IFNγ, and TNF (**Figure 1F**). The presence of IFNγ-related genes, including downstream effector chemokines, support a T helper-1 induced immune response in these myeloid cells. Overall, CD11b+ cells were the most transcriptionally altered in GBS, and the changes were largely unique within CD11b+ and CD8+ cells (**Figure 1G**). There were no differentially expressed genes in CD4+ sorted T cells in the bulk RNA sequencing data.

### Comparative single-cell transcriptomics in AIDP-variant GBS

We next performed single cell-RNA sequencing on unsorted PBMC samples to determine specific, potentially pathogenic leukocyte subpopulation transcriptomic changes in early untreated AIDP-variant GBS. The analysis included 107,656 cells from seven individuals and generated 22 clusters spanning all major PBMC subpopulations (Figure 2A). Cluster identities were manually verified using dimensionality reduction plots (Figure 2B; **Supplemental** Figure 1), heatmaps (Figure 2C), and dot plots (**Supplemental** Figure 2) of cell type-specific marker genes determined by mapping cluster-enriched genes to well-established canonical cell type markers.

**Figure 2.**
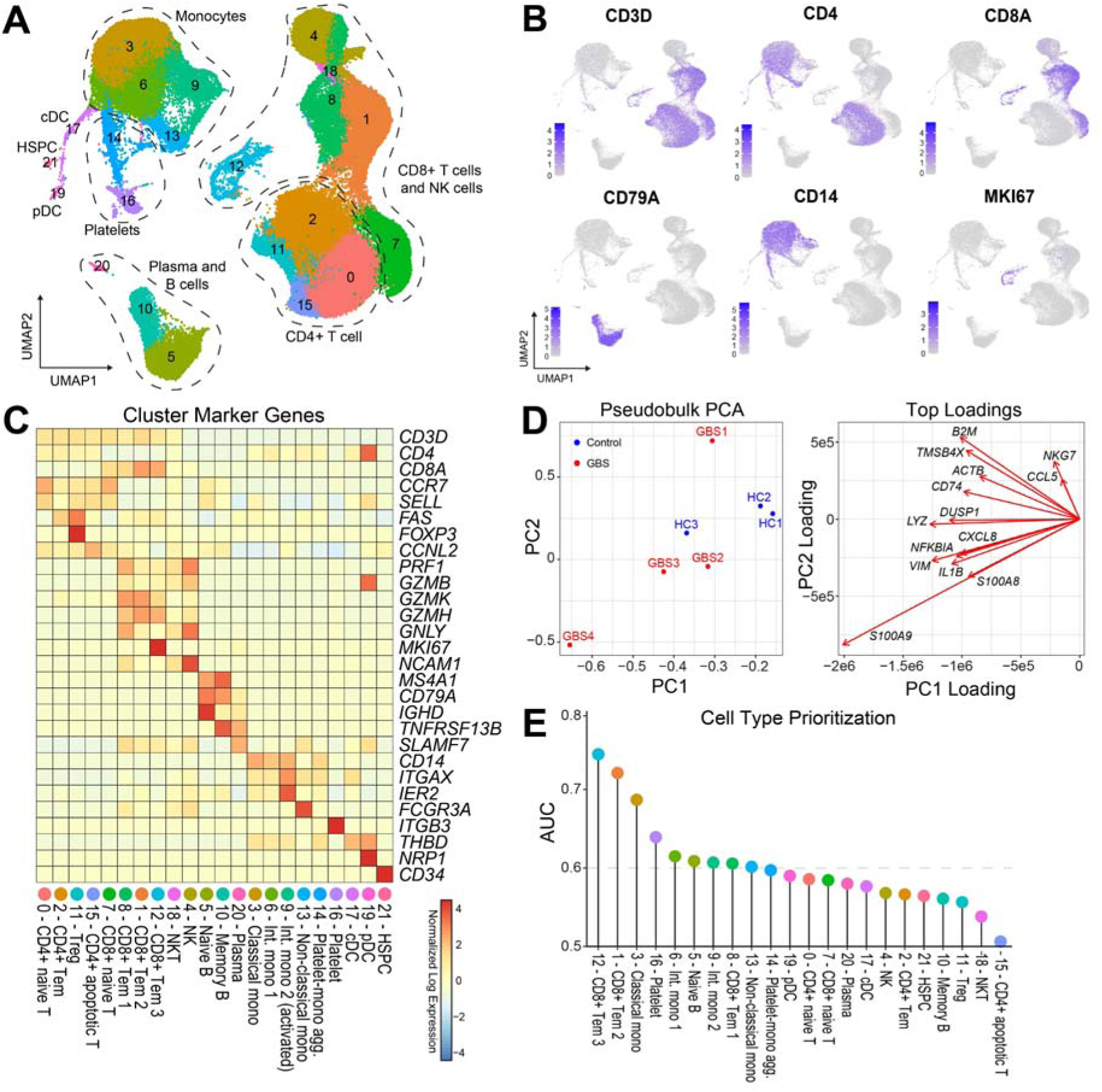
Single-cell RNA sequencing of PBMCs shows gene expression dysregulation associated with early AIDP-variant GBS. **(A)** UMAP dimensionality reduction and Louvain clustering generated 22 unique clusters representing all major peripheral blood leukocyte populations. **(B)** Dimensionality reduction feature plots and **(C)** heatmap of differentiating marker genes used to identify and annotate clusters. **(D)** Principal components analysis (PCA) of pseudobulked gene expression data by sample shows separation of GBS and control samples. Top loadings show the key genes driving separation of samples. **(E)** Augur cell type prioritization found good predictive performance of CD8+ Tem 3, CD8+ Tem 2, and classical monocytes, and some predictive potential from most other cell subpopulations. Dotted line denotes an arbitrary AUC > 0.6 threshold. AUC: area under the curve, where 0.5 denotes a model with performance no better than chance and 1.0 denotes a model with perfect performance in classifying samples into GBS or control based solely on gene expression perturbations within the indicated cluster. cDC: conventional dendritic cell; HSPC: hematopoietic stem and progenitor cells; NK: natural killer cell; NKT: natural killer T cell; pDC: plasmacytoid dendritic cell; Tem: effector memory T cell; Treg: regulatory T cell. cDC: conventional dendritic cell; HSPC: hematopoietic stem and progenitor cells; NK: natural killer cell; NKT: natural killer T cell; pDC: plasmacytoid dendritic cell; Tem: effector memory T cell; Treg: regulatory T cell.

The cell clusters included four *CD3D*+*CD4*+ T cell subsets, consisting of *CCR7+SELL+FAS*lo naïve T cells (cluster 0); *CCR7-SELL-FAS*hi effector memory T cells (Tem; cluster 2); *FOXP3*+ regulatory T cells (Treg; cluster 11); and *CCNL2*-expressing apoptotic-like T cells (cluster 15). There were four *CD3D*+*CD8A+* T cell subsets, including *CCR7+SELL+* naïve T cells (cluster 7) and three cytotoxic T lymphocyte (CTL) subpopulations. *CD8A*+ Tem 1 cells were *PRF1*hi, *GNLY+*, and *GZMH*lo (cluster 8); *CD8A*+ Tem 2 cells were *PRF1*lo*, GNLY-*, and *GZMH*hi (cluster 1); and *CD8A*+ Tem 3 cells had a highly activated Tc1 phenotype and expressed the proliferation marker *MKI67* (cluster 12). The remaining five lymphocyte clusters included *CD3D+NCAM1+* natural killer T (NKT) cells (cluster 18); *CD3D-NCAM1+* NK cells (cluster 4); *IGHD+CD27-* naïve B cells (cluster 5); *TNFRSF13B*+*CD27+* memory B cells (cluster 10); and *SLAMF7*+ plasma cells (cluster 20).

Monocytes were distributed across five clusters: *CD14+FCGR3A-* classical monocytes (cluster 3); *CD14-FCGR3A+ non-*classical monocytes (cluster 13); two *CD14+FCGR3A+* intermediate monocyte clusters, which included one conventional (Int. mono 1; cluster 6) and one highly activated subset with high expression of *ITGAX* and *IER2* (Int. mono 2; cluster 9); and *CD14+ITGB3+* putative platelet-monocyte aggregates (cluster 14). Other clusters included *ITGB3*-expressing platelets (cluster 16); conventional dendritic cells (cDC; cluster 17); plasmacytoid dendritic cells (pDC; cluster 19); and *CD34*+ hematopoietic stem and progenitor cells (HSPC; cluster 21).

To begin exploring differences between the GBS and control cohorts, per-sample principal components analysis (PCA) was performed on pseudobulked gene expression data (Figure 2D). There was good separation between GBS and control samples on the first two principal components, though GBS1 had a more extreme PC2 coefficient compared to the other GBS samples. Plotting the top loadings for PC1 and PC2 showed that PC2 was driven strongly by *NKG7* and *CCL5*. These are both highly expressed in CD8+ T cells and NK cells, suggesting GBS1 has a cellular phenotype driven by cytotoxic lymphocytes. PC1 was instead driven by monocyte genes such as *LYZ* and the cytokine *IL1B*. Notably, the most extreme sample in PC1 and PC2 loadings was GBS4, who presented with the most severe clinical phenotype in this cohort including the highest complete blood count and the highest CSF protein and leukocyte levels. The loadings for this sample were driven by *S100A8* and *S100A9* expression, suggesting a strongly proinflammatory myeloid phenotype. The remaining samples, GBS2 and GBS3, displayed an intermediate phenotype with *LYZ*, *IL1B*, and *NFKBIA* among the contributing loadings.

Next, to help direct the focus of downstream differential analysis, we prioritized cell clusters by training a machine learning model on the gene expression data of each cluster using Augur (28). This generated a sensitivity/specificity area under the curve (AUC) measure of how well GBS or control status can be predicted based solely on gene perturbations within a given cluster (Figure 2E). AUC > 0.6 was used as an arbitrary cut-off. CD8+ Tem 3 proliferating cells (cluster 12) were the most predictive of GBS status (AUC = 0.74), followed by CD8+ Tem 2 (cluster 1; AUC = 0.72). Classical monocytes (cluster 3) also showed reasonable predictive performance (AUC = 0.69). Additional clusters with an AUC > 0.6 included platelets (cluster 16), intermediate monocyte-1 (cluster 6), naïve B cells (cluster 5), highly activated intermediate monocyte-2 (cluster 9), CD8+ Tem 1 (cluster 8), and non-classical monocytes (cluster 13). GBS status in this cohort was therefore best predicted by the phenotype of monocytes and CD8+ Tem cells.

Next, differential abundance analysis was performed to uncover alterations in cell frequency across clusters (Figure 3A). Of the 22 clusters, six were proportionally more abundant in the GBS condition compared to the control condition. The proportion of CD8+ Tem 3 proliferating cells (cluster 12) was starkly higher in GBS blood compared to controls. The remaining five clusters higher in GBS blood were monocyte or platelet clusters: intermediate monocyte-1 (cluster 6), classical monocytes (cluster 3), platelets (cluster 16), platelet-monocyte aggregates (cluster 14), and non-classical monocytes (cluster 13). Six clusters were proportionally lower in GBS, including NKT cells (cluster 18), CD8+ Tem 1 cells (cluster 8), CD4+ apoptotic T cells (cluster 15), memory B cells (cluster 10), pDCs (cluster 19), and NK cells (cluster 4).

**Figure 3.**
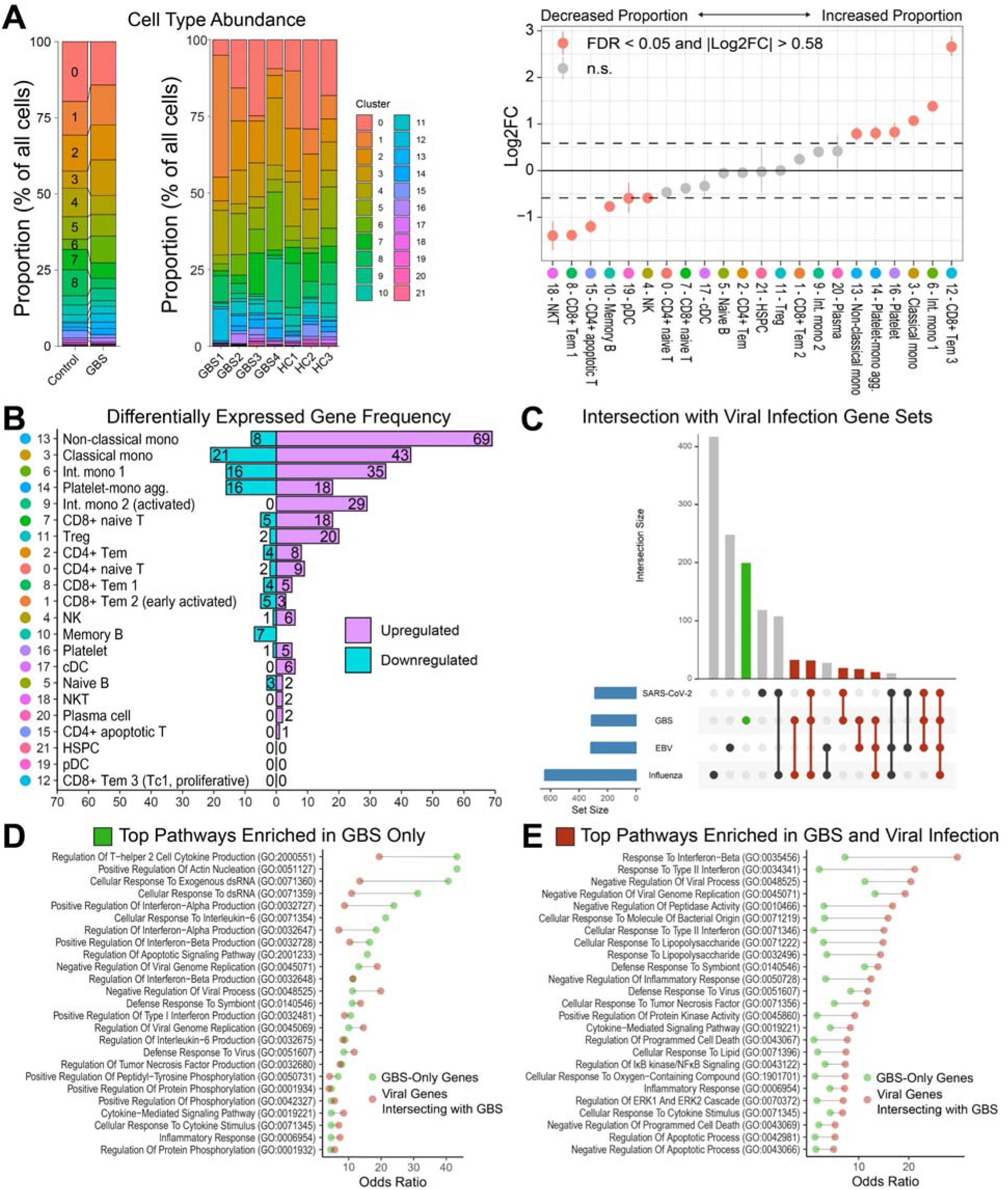
Single-cell RNA sequencing of PBMCs shows gene expression dysregulation associated with early AIDP-variant GBS. **(A)** Differential abundance analysis between GBS and control samples. The stacked bar plots compare the total proportion of cells in each cluster between GBS and control conditions (left) and in individual samples (middle). The permutation plot (right) shows the result of differential abundance testing with significantly enriched and depleted cell types in GBS marked in red. **(B)** Differential expression analysis showing the number of upregulated and downregulated genes in GBS compared to healthy controls for each cluster, ordered by the total number of differentially expressed genes. Analysis was conducted with DESeq2. **(C)** The genes upregulated in GBS samples contain multiple genes that are not significantly upregulated in publicly available PBMC data sets of influenza, SARS-CoV-2, and Epstein-Barr virus infection. Other genes upregulated in GBS also intersect with those upregulated in one or more of these infections. **(D)** Top 25 GO:BP pathways emerging from GBS-upregulated genes that were not present in any of the viral infection data sets. The top pathways were selected by lowest *P*-values (all *P_adj_* < 0.05) and then ordered by the odds ratio of the GBS-only gene set. **(E)** Top 25 GO:BP pathways emerging from GBS-upregulated genes that were present in at least one of the viral infection data sets included for analysis. The top pathways were selected by lowest *P*-values (all *P_adj_* < 0.05) and then ordered by the odds ratio of the shared GBS-viral infection gene set. cDC: conventional dendritic cell; GO:BP: Gene Ontology Biological Process; HSPC: hematopoietic stem and progenitor cells; NK: natural killer cell; NKT: natural killer T cell; pDC: plasmacytoid dendritic cell; Tem: effector memory T cell; Treg: regulatory T cell.

We then performed differential expression analysis using DESeq2, which revealed numerous differentially expressed genes in these clusters (Figure 3B**; Supplemental Table 3**). In line with the bulk RNA sequencing and Augur results, monocyte populations formed the five most differentially regulated cell clusters in GBS compared to healthy controls. The next most differentially expressed subpopulations were naïve *CD8A*+ T cells and regulatory T cells (Treg). Fewer differentially expressed genes were detected in other cell types. No changes were detected in HSPCs, pDCs, or CD8+ Tem 3 proliferating cells. Taken together, monocytes emerged as the most differentially altered cells in early AIDP-variant GBS PBMCs, with further notable differences detected in CD8+ T cells and Tregs.

Given that some of the GBS samples were derived from patients with a known infectious prodrome, we compared the upregulated genes in this cohort with those detected in common viral infections to reveal shared and unique transcriptional programs. This was achieved by extracting upregulated genes from publicly available RNA sequencing and microarray data sets of PBMCS in the context of acute influenza, SARS-CoV-2, and Epstein-Barr virus infection and comparing with the upregulated genes from the GBS samples combined across bulk and single-cell analyses. This showed that the GBS-upregulated genes consisted of both unique genes and those shared with viral infections (Figure 3C). GBS-upregulated genes that did not intersect with the infection data sets, termed GBS-enriched genes, belonged to biological process pathways related to type I interferon production, interleukin-6 (IL-6) production and response, and Th2 cytokine production. Despite not overlapping with the included viral data sets, GBS-enriched genes were also associated with pathways of viral genome replication and viral defense responses (Figure 3D). Genes common to multiple GBS-enriched pathways – and which therefore appeared to drive the pathways unique to the GBS-enriched gene set – included *IL6*, *STAT3*, *GAS6,* and *IL15*. Within the GBS-upregulated genes that were shared with at least one of the viral infection data sets, the top enriched pathways included responses to type I and type II interferons, response to lipopolysaccharide, and regulation of programmed cell death (Figure 3E). GBS samples were therefore more closely associated with type I interferon production, while viral infection samples were more involved in the response to type I and type II interferons. Though these results should be interpreted cautiously given the different library preparation and sequencing techniques used in these studies, and incomplete data on timing of symptom onset, the results appear consistent with an increase in magnitude and persistence of type I interferon production in GBS cases beyond that seen in viral infection alone. This coincides with greater engagement of IL6- and STAT3-mediated signaling pathways in GBS that did not strongly emerge in viral infection cases.

#### Multiple GBS patient monocyte subpopulations express a signature driven by JAK/STAT signaling

We focused our initial investigation of altered transcriptomic programs in GBS leukocytes on monocytes given their high Augur prioritization, increased proportion, and comparatively high number of differentially expressed genes. We first analyzed the shared and unique genes across monocyte subpopulations, finding that most genes were specific to each cluster and cell-type specific alterations in function (Figure 4A). Classical monocytes showed 43 upregulated and 21 downregulated genes in GBS patients (Figure 4B). The most significantly upregulated gene was *JAK3*, with others including *STAT3*, *FAS, GAS6, CLU, and ADM*, while downregulated genes included *MMP9, PLCB1*, and *COLEC12*. The top 5 PANTHER upregulated gene terms were JAK/STAT, Fas, interleukin, epidermal growth factor receptor (EGFR), and platelet-derived growth factor (PDGF) signaling pathways (Figure 4C). In the intermediate monocyte-1 cluster, 35 upregulated and 16 downregulated genes were observed (Figure 4D). JAK/STAT signaling was again the most significantly upregulated pathway, followed by chemokine- and cytokine-mediated inflammation and blood coagulation (Figure 4E).

**Figure 4.**
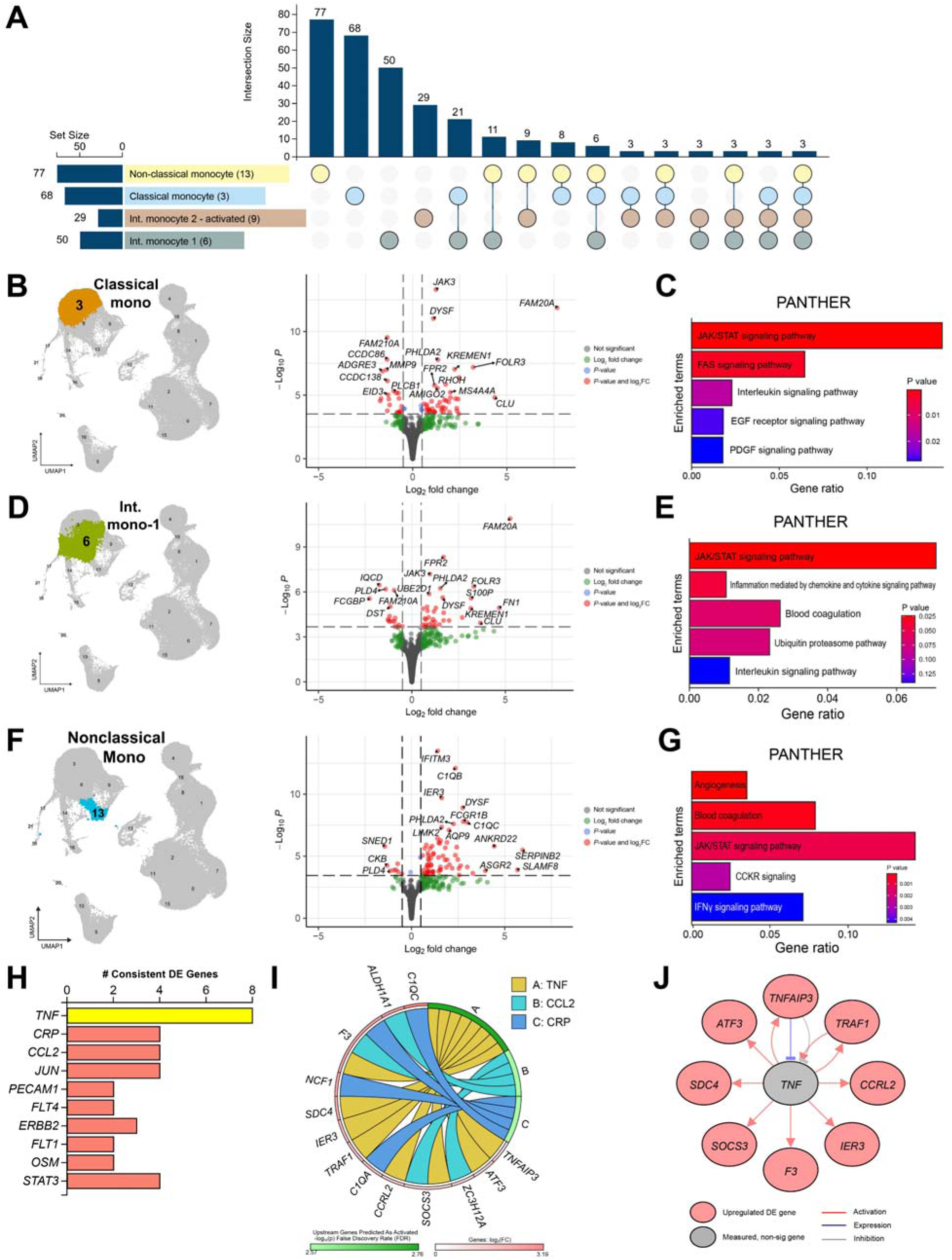
Single-cell transcriptomics of classical (cluster 3), intermediate-1 (cluster 6), and non-classical (cluster 13) monocytes reveals cell-specific responses to shared upstream proinflammatory regulators in GBS. **(A)** Upset plot of differentially expressed genes shared by, or unique to, the four detected monocyte clusters. Non-classical monocytes had the most differentially expressed genes. **(B)** Differential expression analysis of classical monocytes (cluster 3) between GBS and healthy control blood showed 46 significantly upregulated and 23 downregulated genes (*P_adj_* < 0.05 and log_2_FC > |±0.5|). **(C)** Gene set enrichment with PANTHER indicated the most highly associated gene terms in GBS classical monocytes included those associated with the JAK/STAT, Fas, interleukin, EGFR, and PDGF signaling pathways. **(D)** In the intermediate monocyte-1 subset, there were 35 significantly upregulated and 16 downregulated genes in GBS patients. **(E)** The upregulated genes were associated with JAK/STAT signaling, chemokine and cytokine signaling, blood coagulation, the ubiquitin proteasome pathway, and interleukin signaling. **(F)** Non-classical monocytes showed multiple upregulated genes including complement C1q subunits (*C1QA, C1QB, C1QC*), *DYSF*, and Fc-gamma receptor genes (*FCGR1B*). **(G)** The top 5 significantly enriched pathways as determined by PANTHER gene set enrichment analysis of the differentially expressed genes shows angiogenesis, blood coagulation (which included complement-associated genes), JAK/STAT signaling, CCKR signaling, and IFNγ signaling processes. **(H)** Analysis of the top 10 upstream regulators with consistent differentially expressed downstream products shows TNF as the top predicted regulator. TNF, CRP, and CCL2 all reached statistical significance (*P* < 0.05). **(I)** Chord plot shows the differentially expressed genes in non-classical monocytes predicted to be regulated by TNF, CCL2, and CRP. **(J)** The directionality and predicted mechanisms of TNF on differentially expressed genes. CCL2: C-C chemokine ligand 2; CCKR: cholecystokinin receptor; EGFR: epidermal growth factor receptor; IFNγ: interferon-gamma; JAK: Janus kinase; PDGF: platelet-derived growth factor; STAT: signal transducer and activator of transcription; TNF: tumor necrosis factor.

Non-classical monocytes (cluster 13) had the most differentially expressed genes in this cohort, with 69 upregulated and 8 downregulated genes (Figure 4F). Notably, all three complement C1q subunits were upregulated (*C1QA, C1QB, C1QC*), consistent with an enhanced role of these cells in facilitating the initiation of the classical complement pathway directed against antigen-antibody complexes in GBS. This observation was specific to non-classical monocytes, with no C1q genes altered in the other monocyte clusters. The top 5 PANTHER gene terms significantly associated with upregulated non-classical monocyte genes included angiogenesis, JAK/STAT signaling, cholecystokinin receptor (CCKR) signaling, and IFNγ signaling (Figure 4G). The predicted statistically significant upstream regulators of these differentially expressed genes were *TNF*, *CRP*, and *CCL2*, with further nonsignificant regulators in the top 10 most associated including *JUN*, *ERBB2*, *OSM*, and *STAT3* (Figure 4H). The genes driving these upstream regulator predictions included complement C1q genes (*C1QA, C1QC*), *SDC4*, *IER3*, and *ATF3* (Figure 4I-J). Overall, monocyte subsets in GBS upregulated soluble inflammatory factors and associated receptors in a cell-type specific manner in response to shared upstream proinflammatory regulators.

#### A highly activated intermediate monocyte subset is uniquely driven by oncostatin M signaling in early AIDP-variant GBS

In addition to these canonical monocyte subpopulations, a subpopulation of highly activated intermediate monocytes (cluster 9) was observed that was phenotypically distinct from canonical intermediate monocytes (Figure 5A). These highly activated intermediate monocytes expressed genes indicative of strong activation (*IER2*, *ITGAX*) and spliceosome and nucleosome assembly. The gene enrichment terms that most strongly distinguished this subset from the other intermediate monocyte cluster (6) were systemic lupus erythematosus and mRNA surveillance.

**Figure 5.**
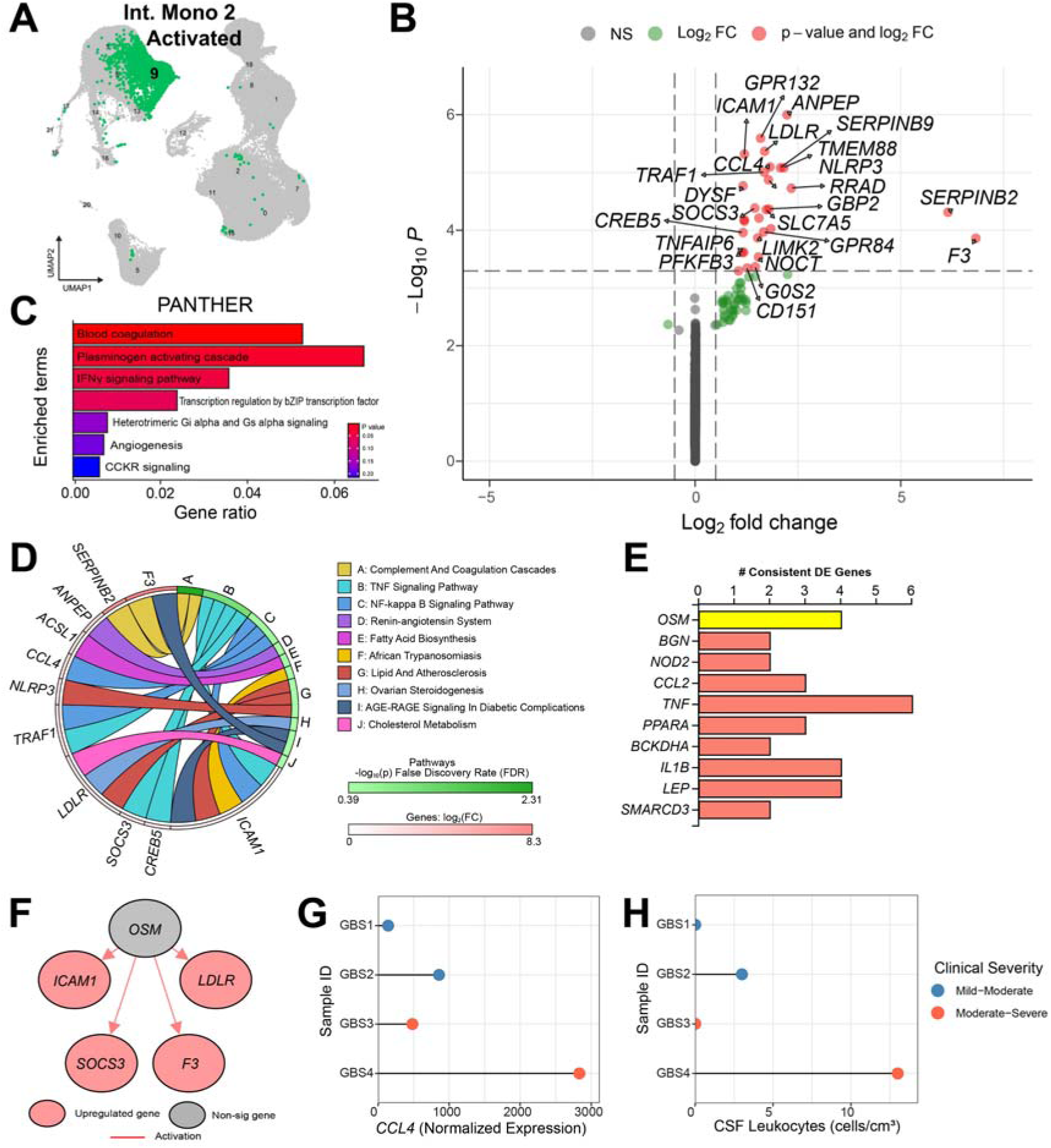
A distinct subpopulation of intermediate monocytes with a highly activated phenotype upregulates genes in GBS predicted to be driven by upstream oncostatin M (OSM) signaling. **(A)** UMAP highlighting the activated intermediate monocytes (cluster 9) in the single-cell data set. **(B)** There were 29 significantly upregulated genes and zero downregulated genes (*P_adj_*< 0.05 and log_2_FC > |±0.5|) within this cluster in GBS patients compared to controls. **(C)** PANTHER gene set enrichment analysis of the significantly upregulated genes of highly activated intermediate monocytes displaying the top 7 pathways. The top three pathways were significantly enriched: blood coagulation (*P* = 0.001), plasminogen activating cascade (*P* = 0.022), and interferon-gamma (IFNγ) signaling (*P* = 0.040). (**D-F**) Pathways were generated and upstream regulators predicted using iPathwayGuide. **(D)** KEGG pathway analysis revealed additional upregulated pathways, including TNF and NFκB signaling. **(E)** The predicted upstream regulators of the differentially expressed genes. The IL-6 receptor family cytokine OSM was the most consistent with the GBS patient upregulated genes. The top 10 statistically significant consistent upstream regulators are depicted (all *P* ≤ 0.005). **(F)** The genes consistent with upregulation by OSM were *ICAM1, SOCS3, F3,* and *LDLR*. In each case, OSM was predicted to activate the gene. **(G)** The normalized expression of *CCL4*, one of the upregulated differentially expressed genes in highly activated intermediate monocytes, in each GBS sample. GBS4, which had one of the most severe physical and electrodiagnostic presentations in this cohort, had the highest level of expression. **(H)** The frequency of leukocytes in the cerebrospinal fluid of each GBS patient. The donor with the highest *CCL4* expression in this cluster also had the highest number of cerebrospinal leukocytes.

These activated intermediate monocytes showed 29 upregulated genes in GBS patients, with select genes including *NLRP3, ICAM1*, and *CCL4* (Figure 5B). PANTHER gene set enrichment identified significant associations with processes of blood coagulation, plasminogen activation, and IFNγ signaling (Figure 5C). Further gene set enrichment by KEGG suggested upregulation of TNF (*ICAM1, SOCS3, CREB5, TRAF1*) and NFκB (*CCL4, ICAM1, TRAF1*; Figure 5D) signaling. These upregulated genes were highly consistent with upstream regulation by the IL-6 family cytokine oncostatin M (OSM). Additional significantly associated upstream regulators included CCL2, TNF, PPARA, IL1B, and LEP (Figure 5E-F).

Given the propensity for activated intermediate monocytes to be recruited to inflammatory sites in tissues, we reasoned that the cluster 9 highly activated intermediate monocytes are among the most likely cells in this data set to extravasate and participate in clinically important interactions in GBS. To explore this possibility, we investigated whether one of the proinflammatory chemokines upregulated in GBS monocytes, *CCL4*, covaried with individual patient clinical information. We found that the patients with the greatest clinical severity, as determined by physical presentation and electrodiagnostics, also had among the highest *CCL4* expression in these monocytes (Figure 5G). The number of leukocytes in the cerebrospinal fluid of these patients was also highest in those with high intermediate monocyte *CCL4* (Figure 5H). Though correlational, this supports the idea that the transcriptomic phenotype observed in these cells may be involved in the clinical manifestations observed in these GBS patients.

### CD8+ T cell subpopulations are JAK/STAT-activated and highly proliferative in GBS

We next focused our investigation on the lymphocyte clusters with the most pronounced alterations in proportion, gene expression, and/or cell type prioritization, which were *CD8A*+ T cells and regulatory T cells. *CD8A*+ T cells clustered into one naïve and three memory subpopulations, with *CD8A*+ Tem 3 cells (cluster 12) significantly enriched in GBS at the cohort level. Per-sample proportion analysis of these four *CD8A*+ T cell subpopulations showed considerable variation between individuals. Nevertheless, *CD8A*+ Tem 3 cells were consistently elevated in GBS samples, including in the GBS1 sample which showed substantial enrichment (Figure 6A). To assist in further ascertaining the identities of the three Tem subpopulations, the enriched genes in one cluster compared to the other two Tem clusters were passed through enrichment analysis with PANTHER. This showed that pathways of apoptosis, TGF-β, angiogenesis, and B cell activation, among others, were enriched in Tem 2 cells (Figure 6B). In contrast, Tem 3 cells were overwhelmingly associated with DNA replication and cell cycle-related pathways, which supports its designation as a cycling and proliferating subpopulation (Figure 6C). Tem 1 cells did not show any robust pathway enrichment and so was considered a basal or canonical *CD8A*+ Tem population.

**Figure 6.**
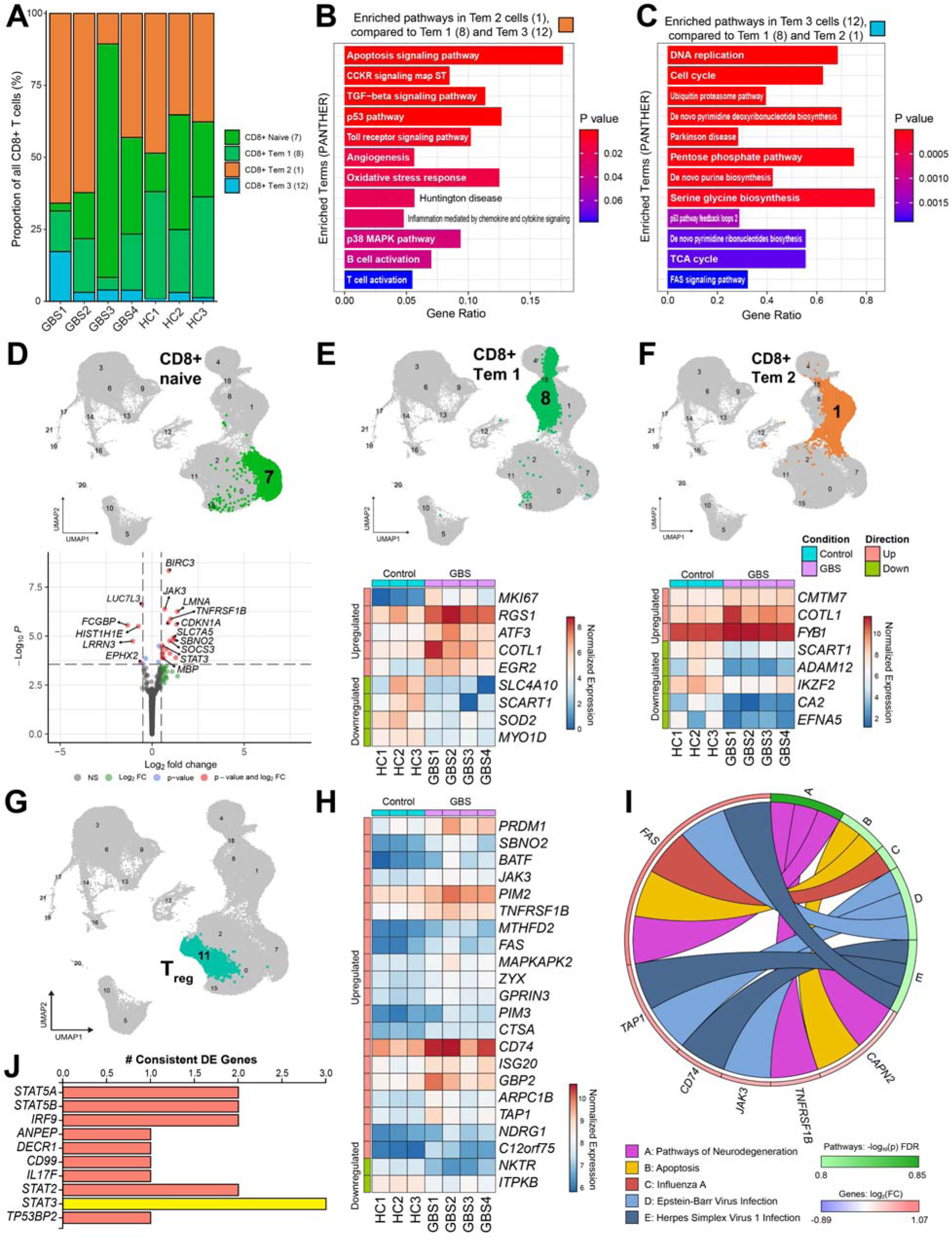
Single-cell transcriptomics of lymphocyte subpopulations shows CD8+ T cell proliferation and regulatory T cell dysfunction in GBS. **(A)** *CD8A+* T cell subpopulation proportions across samples. *CD8A+* Tem 1 cells are decreased in most GBS samples compared to controls, while Tem 3 cells are higher in all GBS samples but particularly in GBS1. **(B)** Gene pathways distinguishing Tem 2 cells from Tem 1 and Tem 3 cells include apoptosis, TGF-β signaling, and lymphocyte activation. **(C)** Gene pathways distinguishing Tem 3 cells from Tem 1 and Tem 2 cells show that these are proliferating *CD8A*+ Tem cells. **(D)** Naïve T cells were the most transcriptionally altered *CD8A*+ T cell subpopulation in GBS, with key upregulated genes including *JAK3*, *STAT3*, *TNFRSF1B*, and *MBP*. **(E)** *CD8A+* Tem 1 cells were less transcriptionally dysregulated but did show upregulated *MKI67* indicative of cell proliferation and *ATF3* associated with cellular stress responses. **(F)** *CD8A+* Tem 2 cells were not substantially altered in GBS, though did significantly downregulate the transcription factor *IZKF2* (Helios) among other genes. **(G)** Differential analysis of regulatory T cells (Treg) shows **(H)** the differential expression of 22 genes (*P_adj_* < 0.05 and log_2_FC > |±0.5|) including *PRDM1* (Blimp-1), *BATF*, *FAS,* and *CD74*. **(I)** Pathways associated with the detected changes in Tregs included viral infection, apoptosis, and pathways of neurodegeneration. Pathways were generated using iPathwayGuide. **(J)** Top 10 statistically significant (all *P* < 0.05) predicted upstream regulators of the differentially expressed genes included multiple STATs. STAT3 was the upstream regulator with the highest predicted log_2_FC (0.681), marked in yellow. Regulators are listed in ascending order by *P*-value.

*CD8A*+ naïve T cells were elevated in genes including *JAK3*, *STAT3*, *TNFRSF1B* (TNFR2), and *MBP* which encodes myelin basic protein (Figure 6D). This may represent an increase in the alternatively spliced Golli-MBP, which has been suggested to play a role in the promotion of tolerance against MBP in nervous system autoimmunity (29, 30), though alternate sequencing methods capable of confident isoform detection would be required to confirm this possibility. *CD8A*+ Tem 1 cells showed upregulation of *MKI67* and *ATF3*, consistent with proliferation and stress response augmentation, respectively (Figure 6E). The few detected changes in *CD8A*+ Tem 2 cells included decreased *SCART1* (encoding the γδ-associated scavenger receptor CD163c-alpha) and transcription factor *IKZF2* (Helios; Figure 6F). Early GBS *CD8A*+ T cells are highly proliferative and undergo increases in JAK/STAT-mediated viral and cell stress responses. However, despite being among the top predictors of GBS status, the gene expression alterations of *CD8A*+ T cells were not as robust as those seen in monocytes in early GBS.

### GBS patient regulatory T cells upregulate transcripts related to persistence and effector function

The lymphocyte subpopulation with the highest number of upregulated genes was regulatory T cells (Tregs), with 20 upregulated and 2 downregulated genes (Figure 6G). The most strongly upregulated gene in Tregs was *PRDM1* (Blimp-1), a marker of effector Tregs and master transcription factor critical for IL-10 production and the persistence of Treg identity in states of high inflammation (31–33). *BATF* expression was also upregulated, which promotes the differentiation of effector and activated Tregs and is important for Treg stability (34–36) but also Th17 differentiation (37). The upregulation of *CD74* was of interest since Tregs with a CD74hi phenotype undergo type I interferon-induced exhaustion in systemic lupus erythematosus (SLE) (38). Given the strong type I interferon response observed in these samples, this raises the possibility that a similar exhaustion occurs in GBS which results in poor Treg effector function. Other notable upregulated genes included *JAK3*, *FAS*, *TNFRSF1B* (TNFR2), *MAPKAPK2*, and *GBP2* (Figure 6H). KEGG pathway analysis indicated enrichment of genes associated with pathways of neurodegeneration, apoptosis, and viral infection (Figure 6I). We found nine statistically significant upstream regulators of these genes, four of which were STATs (*STAT2, STAT3, STAT5A, STAT5B*). *STAT3* had the highest log2 fold-change (0.681) alongside predicted regulation by *IL17F*, which is canonically associated with type 17 immune responses (Figure 6J). These results support a role for effector Treg dysfunction and raise the possibility of exhaustion and identity challenge in early AIDP-variant GBS pathogenesis.

### Monocyte-derived CCL4, epiregulin, and type I interferons are among predicted modulators of GBS Schwann cell and sensory neuron function

Many of the observed upregulated genes in early untreated GBS patient PBMCs encode proteins capable of initiating intercellular transmembrane signaling cascades. We aimed to ascertain which of the genes upregulated in specific GBS patient leukocyte subsets might encode proteins that facilitate direct molecular interactions with peripheral nervous system cells. This enables the prediction of specific molecules expressed by GBS patient leukocytes that are pathologically relevant drivers of peripheral nerve injury. We approached this computationally by mapping the upregulated differentially expressed GBS genes to receptor genes expressed by human dorsal root ganglion (DRG) cells using a previously published and manually curated library of known ligand-receptor interactions (24). We focused our analysis on predicted interactions with myelinating Schwann cells, for their importance in the underlying demyelinating pathology of AIDP-variant GBS, and on sensory neurons to discover pronociceptive ligands that may potentially activate or sensitize them to drive pain.

Hundreds of predicted interactions were observed, which were wholly driven by myeloid cells, confirming these as key effector cells in early GBS. Upregulated genes in the CD11b+ bulk transcriptomics data formed 156 interactions with Schwann cells and 72 interactions with sensory neurons. These interactions were mostly with signaling transmembrane receptors, including G protein-coupled receptors, receptor tyrosine kinases, and serine/threonine protein kinase receptors, as well as those with cell adhesion-related functions. The single-cell data recapitulated shared interactions between monocyte subpopulations but also revealed cell type-specific interactions. Overall, Schwann cells interacted most frequently with classical monocytes, which aligns with our observation that classical monocytes were among the stronger predictors of GBS status. Sensory neurons were instead predicted to interact closely with intermediate monocytes (Figure 7A).

**Figure 7.**
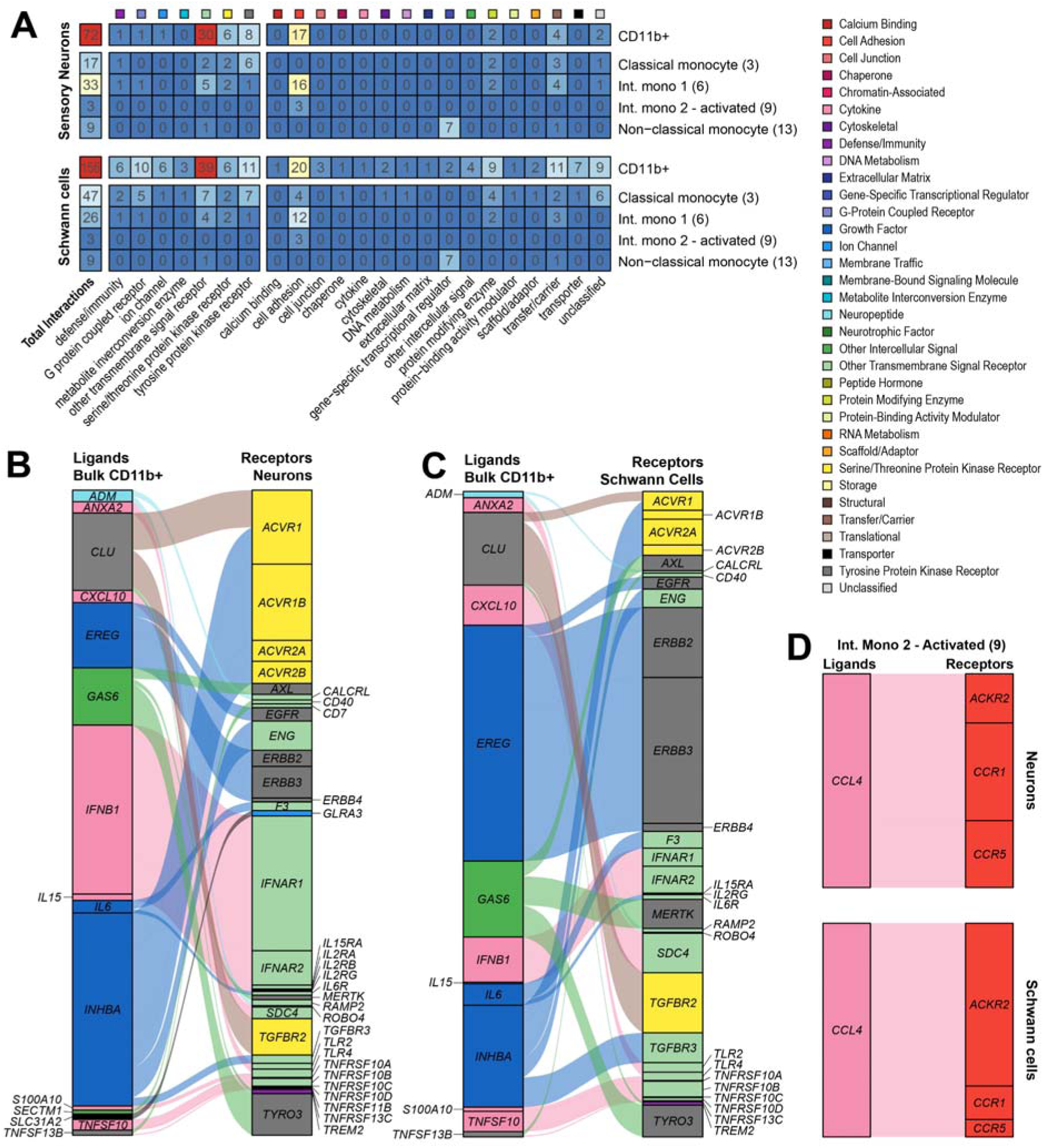
Predicted interactions between upregulated ligand genes in GBS patient PBMCs and receptors expressed in human dorsal root ganglion (DRG) sensory neurons and Schwann cells reveal candidate protein products with possible damage-promoting and/or pronociceptive effects in GBS. **(A)** The number of interactions by myeloid cell type and category of molecular interaction. Sensory neurons had the most predicted interactions with intermediate monocytes, while Schwann cells interacted most with classical monocytes. Most interactions were with transmembrane signaling receptors, including G protein-coupled receptors, tyrosine protein kinase receptors, and other transmembrane receptors. Interactions promoting cell adhesion were also common. **(B)** Ligands corresponding to genes upregulated in the bulk sequencing of GBS patient CD11b+ leukocytes and their predicted interactions with receptors expressed on Schwann cells and **(C)** sensory neurons of the human DRG. Ligands and receptors were colored by their molecular category/function. Alluvial flows of each interaction were colored according to the molecular category of the corresponding ligand and sized by the relative RNA expression (normalized transcripts per million, nTPM) of receptors in the relevant cell type as reported in a publicly available single-nuclei RNA sequencing data set (https://sensoryomics.shinyapps.io/Interactome/). These interactions were limited to those with receptors categorized as belonging to one of the seven molecular classes in the left part of panel A. **(D)** Ligand-receptor interactions between activated intermediate monocytes (cluster 9) and receptors expressed in Schwann cells and sensory neurons reveal CCL4 as a ligand with predicted ability to bind CCR1, CCR5, and ACKR2 receptors on these cells. Colors in all panels denote the molecular classification of ligands and receptors as listed in the figure legend.

The myeloid cell-derived ligands driving Schwann cell and sensory neuron interactions were largely shared, though divergences in receptor expression between these cell types differentially weighted these interactions. The ligands included cytokines and chemokines (*IFNB1, CXCL10, IL6, TNFSF10*/TRAIL*, TNFSF13B*/BAFF, *IL15*), growth factors (*INHBA*, *EREG*) and peptide hormones (*ADM*). In Schwann cells, *EREG* (epiregulin) and its interactions with epidermal growth factor receptor subunits (*EGFR, ERRB2, ERRB3, ERRB4*) were among the most highly weighted. In neurons, the strongest interaction was between *IFNB1* and the two components of the interferon-alpha/beta receptor *IFNAR1* and *IFNAR2*. Type I interferons have been observed to drive pain behaviors in animal models (39) and sensitize human sensory neurons (40). *CLU* (clusterin or apolipoprotein-J) was also predicted to interact with multiple proteins in these cells, which may play a role in nerve regeneration and remyelination (**Figure 7B-C**) (41, 42).

To determine the specific myeloid cell subpopulations with unique contributions to the ligand-receptor interactome in AIDP-variant GBS, we utilized the single-cell data and focused specifically on the ligand genes upregulated in the highly activated intermediate monocyte cluster (9). These cells exhibited a phenotype indicative of preparedness to extravasate, implying that they have a high likelihood of transmigrating from peripheral circulation into peripheral nerves and nerve roots. The three interactions that emerged in both Schwann cells and sensory neurons were driven by the chemokine *CCL4* and directed toward its signaling G protein-coupled receptors *CCR1* and *CCR5* as well as the atypical chemokine receptor *ACKR2* (Figure 7D). Notably, the functional chemokine receptors *CCR1 and CCR5* were the most highly expressed in neurons, while the decoy receptor *ACKR2* was the predominantly expressed receptor in Schwann cells. These interactions were not observed in other monocyte subsets, including the canonical intermediate monocyte cluster (6). This subset instead upregulated *CCL20* which interacted with *CXCR3* expressed on neurons but not on Schwann cells (**Supplemental** Figure 3). Taken together, the transcriptomic changes occurring in untreated AIDP-variant GBS PBMCs are predicted to involve unique molecular interactions. These predictions may be investigated experimentally for their potential contributions to GBS pathogenesis and the generation of associated neuropathic pain.

### CCL4 sensitizes human sensory neurons

We verified the predicted interaction between activated intermediate monocyte-derived CCL4 and neurons by measuring the responses of primary human sensory neurons to recombinant CCL4 using calcium imaging (Figure 8A-B). Pretreatment with 100 ng/mL CCL4 for 30 min significantly increased the proportion of sensory neurons responsive to capsaicin (20 nM) when compared with vehicle-treated neurons, indicating a sensitizing effect of CCL4 (Figure 8C). We further investigated the properties of responses in capsaicin-responsive and unresponsive neurons. The subset of neurons that responded to capsaicin did not show an increased magnitude of response to CCL4 (Figure 8D), showed a non-significant increase in the area under the curve of the response (*t_167.3_*= 1.90, *P* = 0.059; Figure 8E) and a non-significant decrease in the latency of response (*t_166.4_* = 1.90, *P* = 0.059). In capsaicin-unresponsive neurons, CCL4 pretreatment did not significantly increase baseline intracellular calcium (Figure 8F) but did significantly increase their responses to KCl, suggesting an overall increase in the excitability of these sensory neurons (Figure 8G). The baseline calcium in capsaicin-responsive neurons was no different between CCL4- and vehicle-treated neurons (*t_168.9_* = 0.006, P *>* 0.99), which provides evidence against the induction of spontaneous activity following 30 min exposure to CCL4. These results confirm the predicted ability of CCL4 to sensitize human sensory neurons and provide support for the pronociceptive potential of mediators released by activated monocytes in AIDP-GBS.

**Figure 8.**
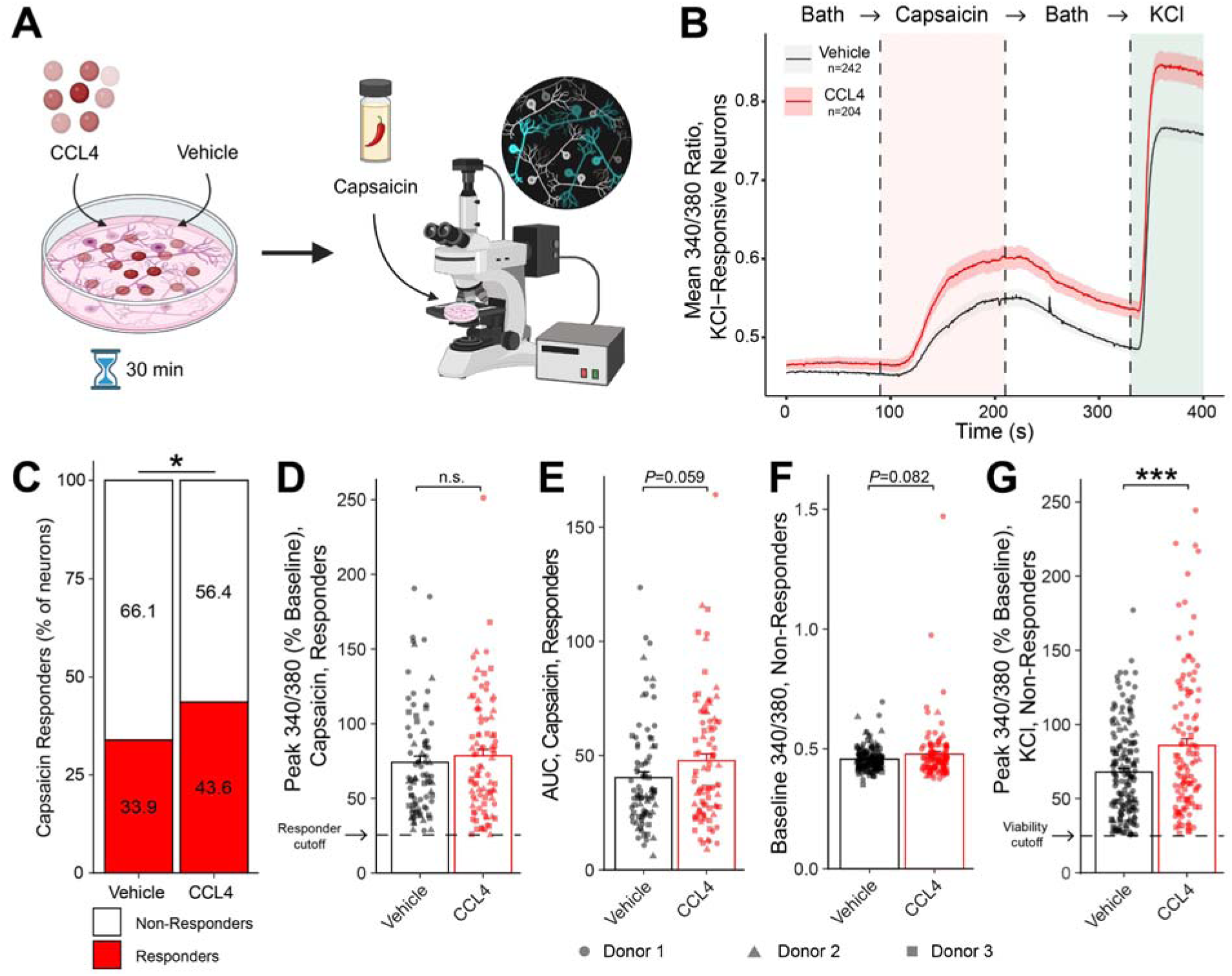
CCL4 sensitizes primary human sensory neurons *in vitro* as measured by calcium imaging. **(A)** Dissociated primary human DRG neurons from three donors were pre-treated for 30 min with 100 ng/mL CCL4 or vehicle. Cells were then removed from the pretreatment medium and underwent calcium imaging in response to the TRPV1 agonist capsaicin (20 nM) and potassium chloride (KCl, 50 mM). **(B)** Mean calcium traces for CCL4- and vehicle-treated neurons across all donors. CCL4 pretreatment slightly increased basal intracellular calcium, robustly increased responses to capsaicin, and additionally increased KCl responses. The *n* values refer to the number of cells per condition. **(C)** Significantly more neurons were responsive to capsaicin (>25% increase in 340/380 ratio) when pretreated with CCL4 compared to vehicle. Fisher’s exact test, *P* = 0.040. **(D)** The magnitude of response to capsaicin within capsaicin-responsive neurons was not significantly altered after CCL4 pretreatment (Welch’s *t*-test, *t_168.9_* = 0.75, *P* = 0.451). **(E)** The area under the curve (AUC) of the capsaicin response was not significantly elevated in capsaicin responders. **(F)** The baseline 340/380 ratio was not significantly different in CCL4-treated capsaicin-unresponsive neurons compared to vehicle neurons (*t_139.7_* = 1.75, *P* = 0.082). **(G)** The KCl response magnitude was significantly increased in CCL4-treated capsaicin-unresponsive neurons (*t_182.6_* = 3.52, *P* = 0.0005). \**P* < 0.05, \*\*\**P* < 0.001. Panel A was created with BioRender.com.

## Discussion

Monocytes and T cells were the principal transcriptionally dysregulated leukocyte subpopulations in this cohort of early AIDP-variant GBS patients. Janus kinase (JAK) and signal transducer and activator of transcription (STAT) signaling, OSM-associated gene expression, and the proliferation of cytotoxic T cells were among the key processes driving this dysregulation. Despite no changes in bulk CD4+ cells, numerous changes were detected specifically in Tregs consistent with dysfunction and potential exhaustion. Classical, intermediate, and non-classical monocytes upregulated genes encoding proteins that may interact with peripheral Schwann cells and sensory neurons via ligand-receptor pairs, potentially contributing to demyelination, axonal degeneration, and/or pain. Finally, CCL4 was identified as a potentially pronociceptive mediator in GBS, since it is sufficient to sensitize primary human sensory neurons *in vitro*.

Based on our findings, JAK/STAT pathways, including JAK3, STAT1, and STAT3, are among the primary transcriptional drivers of the early AIDP-variant GBS immune response. Previous studies have identified monocytes as a key factor in the GBS immune response with the involvement of IL-1β, transforming growth factor (TGF)-β, and CCL2 signaling, which we recapitulated in the current study, but JAK/STAT signaling had not previously emerged (43). This could speak to the heterogeneity in patient disease duration in the cohort of this previous study and may suggest that JAK/STAT signaling is specific to the early and/or untreated phase. CD8+ T cells were highly proliferative in GBS patients and displayed altered expression levels of several transcription factors, including JAK3, STAT3, and Helios. Our results in CD8+ T cells suggest increased capability for cytotoxic responses. This is in line with another study which identified type 1-activated T cells engaging in cytotoxic processes more so than those obtained from influenza patients in GBS patient blood-derived leukocytes (9).

Treg dysregulation may be hypothesized to result in a failure of peripheral tolerance, enabling an inappropriate immune response against myelin antigens. Treg frequency is reduced in early GBS while Th17 cells expand (21, 22, 44–46), an effect reversed by intravenous immunoglobulin treatment (46–48). In our data set, the most highly upregulated gene in GBS Tregs was *PRDM1*, encoding Blimp-1, which is a master transcriptional regulator that maintains Treg identity in the face of inflammatory challenge (31) and is thought to mark an antigen-experienced effector memory phenotype (32). The phenotype observed here, which is high in both *PRDM1* and the MHCII chaperone *CD74* (49), is similar to a subset of Tregs identified in SLE patients that have been functionally exhausted by type I interferon overstimulation (38). Functional studies investigating the response properties of early stage AIDP-variant GBS patient-derived Tregs may provide further characterization of the putative role of Treg dysregulation in tissue-specific autoimmunity.

### Monocyte-Schwann cell interactions suggest possible pathways in demyelination

We identified a set of *GAS6* interactions with *TYRO3*, *AXL,* and *MERTK* (the so-called TAM receptors) that link AIDP-variant GBS patient monocytes to Schwann cells. Gas6 is elevated in the cerebrospinal fluid of chronic inflammatory demyelinating polyneuropathy patients (50). Gas6 to Axl signaling promotes the clearance of myelin debris by macrophages after nerve injury, and Schwann cells upregulate their expression of TAM receptors following injury to facilitate phagocytosis (51). This raises the question of whether peripheral nerve-infiltrating leukocytes in GBS induce Schwann cell TAM upregulation facilitating inappropriate phagocytosis by highly activated transmigrated intermediate monocytes. Other monocyte-Schwann cell interactions included *TNFRSF10* (TRAIL), which may directly interact with Schwann cell receptors to promote apoptosis (52), and *IL15,* which induces JAK/STAT signaling via its receptor subunits *IL15RA* and *IL2RG* (53). A potential pathogenic role of IL15 in Schwann cells has not yet been explored.

### Epiregulin, oncostatin M, and CCL4 are potential mediators of nociception in GBS

Many of the neuroimmune ligand-receptor interactions emerging from this analysis are known pain-promoting pathways. Type I interferons modulate pain in the peripheral and central nervous systems (40, 54–56) and directly interact with interferon receptors on nociceptors to drive inflammatory (57) and neuropathic pain (39).

Epiregulin (*EREG*), an endogenous ligand of the epidermal growth factor receptor (EGFR), was upregulated in myeloid cells. EREG was the top-weighted GBS-associated ligand signaling to myelinating Schwann cells, which express EGFR (58). The functional effects of EREG-Schwann cell signaling are unknown; however, EREG-EGFR interactions drive inflammatory fibrosis (59), so this interaction in peripheral nerves may contribute to persistent Schwann cell injury, demyelination, and axonal degeneration in GBS. Interactions between EREG and sensory neurons are pronociceptive (60), implying these circulating myeloid cells may interact with sensory neurons to mediate nociception in GBS. Modulating myeloid cell EREG might represent a therapeutic target for GBS and neuropathic pain relief, provided future studies support or validate our hypothesized role of EREG-EGFR signaling in GBS pathogenesis. The EREG- and TGF-α-targeting antibody Fepixnebart is currently undergoing clinical trials for diabetic neuropathic pain, chronic low back pain, and osteoarthritis (61, 62), providing proof of concept that may translate to GBS.

Our study predicted that oncostatin M (OSM) is an upstream driver of multiple differentially expressed genes in activated intermediate monocytes in our GBS patient cohort. OSM induces neuronal MAPK signaling to promote nociception in rodents (63, 64). Stimulation of primary human DRG nociceptors with OSM *in vitro* also generates functional and transcriptomic changes associated with chronic neuropathic pain (65). OSM is a potential therapeutic target for chronic neuropathic pain that occurs in GBS patients. GSK2330811, an OSM neutralizing antibody, reported negative phase 2 results for systemic sclerosis (66). However, Vixarelimab, a human monoclonal antibody directed against the OSM receptor beta subunit, demonstrated favorable results in a phase 2b trial for pruritis (67). Small molecule inhibitors of OSM signaling are also being developed (68).

We demonstrated that CCL4 (or MIP-1β) sensitizes human sensory neurons *in vitro*. This supports the notion that CCL4 secreted by activated intermediate monocytes with high transmigration potential in our GBS patient cohort may be pronociceptive. A previous report demonstrated that *CCL4* upregulation is more pronounced in GBS patient CD4+ memory T cells when stimulated *in vitro* with PNS myelin antigens than with influenza antigen, supporting a specific pathogenic role for this chemokine in GBS (9). Preclinical models of neuropathic pain support pronociceptive roles for macrophage- and Schwann cell-derived CCL4 (69, 70). CCL4 binds multiple chemokine receptors, including CCR1 and CCR5. CCR5 is expressed on both neurons and Schwann cells in the human DRG (58), but also other cell types, most notably on resident CD4+ T cells in the human DRG (58, 71). Chemokine ligand-receptor redundancy and promiscuity, expression on multiple cell types under basal physiologic or pathophysiologic conditions, and challenges in chemokine receptor therapeutic modulation have resulted in multiple failed clinical trials over the past 25 years (72–74). Although CCL4 may contribute to nociception in GBS, therapeutically modulating its signaling using direct receptor antagonists may therefore prove challenging.

### Limitations

Our study has important limitations. The cohort size is relatively small, and patient histories (both with and without infectious exposures) may not reflect the full breadth and incidence of GBS presentations present in the broader population. Due to the number of samples tested, it is possible that rare pathogenic PBMC subpopulations or characteristics were not identified. Though many of the key findings were observed in those with and without reported infectious prodromes and were distinct from those observed in common infections, we cannot completely rule out the possibility that some of the observed changes are generic reflections of infection rather than specific drivers of GBS autoimmunity. Future studies with a longitudinal design may be able to shed further light on this possibility. Many of the reported intercellular interactions are based solely on computational predictions and would benefit from further experimental validation. They also require leukocytes to transmigrate across the blood-nerve barrier to participate in their predicted functional effects. This is very likely based on their high activation state and expression of known leukocyte trafficking and cell adhesion transcripts, but identification of these cell types in AIDP-variant GBS patient nerve biopsies would be required to confirm these interactions.

### Conclusions

Monocytes and T cells are the principal dysregulated PBMCs in this cohort of early phase AIDP-variant GBS patients. Transcriptomic pathways enriched in GBS included those associated with JAK/STAT signaling, type I and II interferon activation, and Treg dysfunction. Differential expression data predicted increased molecular interactions between monocytes and both Schwann cells and sensory neurons. This included the chemokine CCL4, which is sufficient to sensitize human sensory neurons to TRPV1 stimulation *in vitro*. The results discussed here provide insight into the peripheral immune environment in the early stages of GBS. The intercellular molecular pathways uncovered by these analyses may represent targets for therapeutic intervention using existing and novel therapies with the aim of preventing, minimizing, or reversing GBS-associated nerve damage and neuropathic pain.

## Methods

### Patient and sample information

Adult patients with AIDP-variant GBS (*n* = 4, 2 female) and age-matched healthy controls (*n* = 4, 2 female) were included in this study. The GBS group consisted of consecutive untreated patients with clinical, electrophysiological and supportive cerebrospinal fluid analytical data consistent with AIDP-variant GBS who presented to a single academic center’s emergency room and consented to give blood for the study.

AIDP-variant GBS was diagnosed by a board-certified academic neuromuscular disease specialist (E.E.U.) based on the revised National Institutes of Neurological Disorders and Stroke (NINDS) criteria (75) with electrodiagnostic characterization as AIDP based on the published Plasma Exchange/Sandoglobulin Guillain-Barré Syndrome Trial Group criteria (76). Patients were at least moderately affected with quadriparesis, distal > proximal > generalized paresthesia or sensory loss, areflexia and ataxia requiring hospitalization, and were eventually treated with intravenous immunoglobulins or plasma exchange. For healthy controls, exclusion criteria included the presence of any neurological disorders, recent or current systemic illness, or infectious exposures.

### Blood sample processing

All GBS patient blood draws were performed within 14 days of reported symptom onset and prior to immunomodulatory treatment initiation. Whole heparinized blood was obtained from GBS patients and controls by a trained phlebotomist, and density gradient centrifugation was immediately performed to isolate PBMCs, followed by controlled rate cryopreservation as previously published (77–80). Cryopreserved leukocytes were stored long-term in liquid nitrogen.

Frozen cryovials containing 10^7^ PBMCs/mL were thawed in a 37°C water bath for 2 minutes. Following thawing, 1 mL of pre-warmed RPMI 1640 medium (Gibco, 11875-093) supplemented with 10% HyClone™ Fetal Bovine Serum (Thermo Fisher Scientific, SH3008803IR) and 1% penicillin-streptomycin (Thermo Fisher Scientific, 15070063) was added to each vial. The cell suspension was transferred to a 15 mL conical tube, and the cryovials were rinsed with additional RPMI media to maximize PBMC recovery. To aid in cell dissociation, Accumax (Thermo Fisher Scientific, 00-4666-56) was added at a 1:1 ratio to the sample volume, and cells were incubated at room temperature for 5 minutes. Samples were centrifuged at 350*g* for 5 minutes at room temperature. The supernatant was aspirated, and the cell pellets were resuspended in 1 mL of RPMI media containing 100 µg/mL DNase I and incubated at 37°C for 15 minutes. At this stage, 350 µL of cells were taken for single cell RNA sequencing and the remainder were passed on for cell sorting and bulk RNA sequencing. One female control (HC4, sample #6) was later excluded due to having a low proportion (<70%) of live cells in the blood sample as determined by flow cytometry viability staining (**Supplemental** Figure 4), resulting in a final sample size of *n* = 4 GBS patients and *n* = 3 controls.

### Cell sorting and bulk RNA sequencing

#### Fluorescence activated cell sorting

Cells were washed with 10 mL of flow cytometry staining buffer (Invitrogen, 00-4222-26) and centrifuged at 350g for 5 minutes at room temperature. Cells were resuspended in 1X phosphate-buffered saline (PBS) and prepared for fluorescence-activated cell sorting (FACS). Cells were incubated with Zombie UV viability stain for 10 minutes at room temperature. Then, 1 mL of flow cytometry staining buffer was added to each tube. Cells were centrifuged, resuspended in flow cytometry staining buffer, and treated with human Fc receptor blocker for 10 minutes at room temperature. The cells were then treated with primary antibodies targeting CD45, CD3, CD4, CD8, and CD11b for 30 minutes, on ice, protected from light (see **Supplemental Table 4** for reagent details). Cells were then washed again with 1 mL of flow cytometry staining buffer, centrifuged, resuspended in flow buffer, and preserved on ice until sorting. FACS was performed on a BD FACSAria Fusion at the University of Texas at Dallas. Cells were gated to collect CD4+, CD8+, and CD11b+ cells in flow cytometry staining buffer (see **Supplemental** Figure 4 for gating strategy).

#### RNA extraction

Tubes containing sorted cells were centrifuged, resuspended in 1 mL of Qiazol (Qiagen, 1023537), transferred to a 1.5 mL Eppendorf tube, and proceeded to RNA extraction. Next, 200 µL of chloroform (Fisher Scientific, C298-500) was added to each tube, manually shaken for 15 seconds, and incubated at room temperature for 2 minutes. Tubes were then centrifuged at 12,000*g* for 10 minutes at 4°C. The supernatant was collected, placed in a new 1.5 mL tube, and 500 µL of isopropanol (Fisher Scientific, A416-1) and 3 µg of GlycoBlue™ (Thermo Fisher Scientific, AM9515) were added. To maximize RNA precipitation, tubes were incubated overnight at -20°C. The next day, the cells were centrifuged at 12,000*g* for 30 minutes at 4°C. The supernatant was aspirated, and the resultant RNA pellet was washed with 1 mL of 75% ethanol (Fisher Scientific, BP2818-500). The tube was centrifuged at 7,500*g* for 5 minutes at 4°C and the supernatant was removed. The resultant RNA pellet was air dried for 10 minutes at room temperature and eluted in 20 µL of RNase-free water on a heat block at 55°C for 10 minutes. RNA concentration and quality numbers were measured on a NanoDrop and the RNA was stored at -80°C until RNA sequencing.

#### Bulk RNA sequencing

RNA integrity numbers (RIN) were assessed and only samples with RIN ≥ 8.0 were used for bulk RNA sequencing (**Supplemental Table 5**). Sequencing was performed using a NextSeq2000 for single-end total RNA sequencing at the University of Texas at Dallas Genome Core Research Facility.

Raw FASTQ files were quality-controlled using Phred scores, per-base sequence, and duplication levels with FastQC (v0.11.0). Reads were soft-clipped (12 bases per read) to remove adapters and low-quality bases, then aligned and sorted using STAR (v2.7.6) (81) with the GRCh38 human reference genome (GENCODE release 38, primary assembly). Sambamba (v0.8.2) (82) was used for deduplication. Prior to quantification, ENCODE blacklist genes were discarded using bedtools (v2.30.0). Data were then passed through StringTie (v2.2.1) to obtain normalized transcripts per million (TPM) count values for each gene (83–85). The Rsubread (v2.14.2) featureCounts function was used to calculate raw counts for all genes. Differential expression analysis was conducted on the resulting output with DESeq2 (86, 87). Genes were considered significantly differentially expressed if *P_adj_* < 0.05 and log_2_ fold-change (FC) > |±0.5|.

### Single-cell RNA sequencing

Cells were washed once with 1X PBS and centrifuged at 500*g* for 5 minutes. The resulting cell pellets were fixed at 4 °C for 17 hours using the Chromium Single Cell Fixed RNA Sample Preparation Kit (10x Genomics). Fixed cells were subsequently hybridized with probes using the Fixed RNA Feature Barcode Kit (10x Genomics), following the manufacturer’s protocol. The remaining library preparation steps were carried out according to the standard 10x Genomics workflow. Libraries were sequenced on an Illumina NextSeq 2000 platform at the University of Texas at Dallas Genome Core Facility. Sequencing data were processed and aligned to the GRCh38 human genome reference using Cell Ranger v7 (10x Genomics).

Individual processed .h5 files corresponding to each sample were imported into Seurat v5.2.1 (88), normalized, and filtered for high-quality nuclei based on the number of features, counts, and percentage of mitochondrial genes (**Supplemental** Figure 5). The final per-sample cell count is reported in **Supplemental Table 6**. Samples were integrated using the canonical correlation analysis (CCA) method of the FindIntegrationAnchors() function. Data were scaled and principal components analysis (PCA) was performed on the top 2000 variable features. Uniform manifold approximation and projection (UMAP) and Louvain clustering (resolution = 0.6) were performed on the top 25 principal components. Cell types were identified and annotated by computing the topmost differentially expressed markers in each cluster, followed by manual verification of cell-specific marker expression using dotplot, heatmap, UMAP, and ridgeline plots.

### Cell type prioritization and differential analysis

Per-sample principal components analysis was prepared by grouping samples and pseudobulking gene expression across all clusters using Seurat’s AggregateExpression() function and performed using the in-built prcomp() function. Cell type prioritization was conducted using Augur v1.0.3 (28). The machine learning model AUC for each cluster was calculated using the calculate_auc() function and visualized with ggplot2 v3.5.2.

Differential abundance analysis between GBS and control samples was performed using scProportionTest v1.0.0 (89). The permutation_test() function was used to find the significance of the log_2_ proportional difference between the two groups. Multiple comparisons correction was performed using a false discovery rate < 0.05 and the 95% confidence interval determined by bootstrapping.

Differential expression analysis was conducted by generating pseudobulk samples from each cluster and comparing GBS to control samples using DESeq2 v1.44.0 (86) with log-fold change shrinkage using apeglm v1.26.1 (90). Strict criteria were used to determine significantly different upregulated and downregulated genes (*P_adj_* < 0.05 and log_2_ fold-change (FC) > |±0.5|) to minimize type 1 error. All *n* used for statistical analysis of scRNA-seq data refer to individual donors and not the number of cells.

### Pathway and gene set enrichment analysis

Gene set enrichment analysis (GSEA) was conducted using enrichR v3.2 (91). Significantly upregulated genes both from cell-sorted bulk-RNA sequencing data and individual scRNA-seq clusters were analyzed for overrepresentation against terms defined by the Gene Ontology (GO) molecular function and biological process (2023 version), KEGG (2016), and Panther (2016) databases. *P* values were generated to determine significantly enriched terms. For plotting, the gene ratio was used as the x-axis label, defined as the overlap between the upregulated gene list and the term-related genes divided by the total number of upregulated genes.

Additional pathway and upstream regulator analyses of both bulk- and scRNA-seq data were performed using iPathwayGuide (Advaita Corporation, Ann Arbor, MI) using its proprietary ‘impact analysis’ to examine pathways, Gene Ontology (GO) terms, and upstream regulatory genes (https://advaitabio.com/ipathwayguide). Pathway analysis examines for DEGs known to be enriched in certain biological pathways; Gene Ontology terms categorize the role of genes in terms of biological processes and molecular function; and upstream regulators are transcripts that are predicted as being present due to their differentially expressed response genes. This approach considers the direction of all signals on a biological pathway along with the location, function, and type of each gene (92–94). Similarities and differences between the bulk CD11b+ sequencing and specific monocyte subsets were elucidated using iPathwayGuide’s meta-analysis feature. All iPathwayGuide analysis underwent multiple comparisons correction for false discovery rate (FDR). Figures using iPathwayGuide outputs are indicated in the corresponding figure legend.

### Comparison with publicly available viral infection data sets

The similarity between the genes upregulated in GBS and those upregulated in a general infectious and inflammatory environment were assessed using pathway analysis on intersected gene sets from previously published data sets from RNA sequencing or microarrays of influenza and SARS-CoV-2 (95) and Epstein-Barr virus (96). Gene sets of upregulated genes in the infectious states reported in each paper were extracted from the supplemental data and intersected with the total list of differentially upregulated genes from both bulk and single-cell RNA sequencing reported in the current results. This generated a ‘GBS-only’ gene set, which included those genes that were upregulated in the present study but not in any of the three infection states, and a shared gene set that were common between GBS and at least one infection state. Upset plots were generated with UpSetR (v1.4.1). Pathway analysis on the GBS-only and shared gene sets was conducted in enrichR (v3.2) using the GO Biological Process 2023 database. The top 25 pathways for each gene set according to adjusted *P*-value (all *P_adj_* < 0.05) were extracted, ordered by the odds ratio in one gene set, and plotted against the odds ratio in the other gene set using Cleveland dot pots in ggplot (v3.5.2).

### Ligand-receptor interaction analysis

Statistically significantly upregulated genes from the bulk-seq data and each scRNA-seq cluster were assembled into lists of candidate ligands. We then intersected this list with a manually curated, publicly available list of experimentally documented ligand-receptor interactions (https://sensoryomics.shinyapps.io/Interactome/; version as of May 2025).

The platform was designed for the identification of physiologically relevant intercellular interactions with the peripheral nervous system, with a particular emphasis on pain targets (24). The list of interactions was filtered based on the following criteria: the ligand was differentially expressed in GBS with P_adj_ < 0.05 and log_2_FC > 0.5; and the receptors for these ligands must be expressed in either sensory neurons or myelinating Schwann cells at a level greater than or equal to 50 normalized transcripts per million (nTPM) per the human DRG single-nuclei and spatial RNA sequencing data set presented in the interactome platform and published elsewhere (25, 26). Interactions were then ranked in descending order based on their receptor counts in sensory neurons/myelinating Schwann cells as appropriate. These interactions were visualized with alluvial plots using *R* packages ggplot2 (v3.5.2) and ggalluvial (v0.12.5).

### Primary human sensory neuron recovery and culturing

Human tissue procurement procedures were approved by the University of Texas at Dallas Institutional Review Board (protocol number: Legacy-MR-15-237) with policies on donor screening and consent provided by the United Network for Organ Sharing.

Human DRGs were recovered from organ donors through the Southwest Transplant Alliance (Dallas, TX) within 4 hours of cross-clamp as described previously (97, 98). Neurons from seven DRGs were used from three organ donors. The DRGs were placed in artificial cerebrospinal fluid for immediate processing or in Hibernate-A medium (Fisher Scientific, NC0176976) supplemented with 1% penicillin/streptomycin (Thermo Fisher Scientific, 15070063), 1% Glutamax (Thermo Scientific, 35050061), 1% N2 (Thermo Scientific, 17502048), 2% NeuroCult SM1 (Stemcell technologies, 05711), 2mM sodium pyruvate (Gibco, 11360-070), and 0.1% bovine serum albumin (BSA; Biopharm, 71-040) for 11.5 hours at 4°C before processing (99). Demographic and sample information is summarized in **Supplemental Table 7.**

DRGs were cultured as previously described (100). DRG bulbs were trimmed, cut into 3mm pieces, and placed in 5mL pre-warmed digestion enzyme containing 1mg/mL Stemxyme I (Worthington Biochemical, LS004106), 0.1mg/mL DNAse I (Worthington Biochemical, LS002139), and 10ng/mL recombinant human β-NGF (R&D Systems, 256-GF) in HBSS without calcium and magnesium (Thermo Scientific, 14170-112).

DRGs were dissociated in a 37°C shaking water bath with trituration every hour until cells were dissociated (∼4.5 hours). The resulting cell suspension was filtered through a 100µm cell strainer (Corning, 431752) and passed through a gradient of 10% BSA in HBSS by overlayering the suspension and centrifuging at 900g for 5 min at RT. The cell pellet was resuspended in media (BrainPhys media, Stemcell Technologies, 05790) containing 1% penicillin/streptomycin, 2% NeuroCult SM1, 1% Glutamax, 1% N2, 10ng/mL β-NGF, 2% fetal bovine serum (Thermo Fisher Scientific, SH3008803IR), 0.1% of 3mg/mL 5-Fluoro-2′-deoxyuridine (Sigma-Aldrich, F0503) and 7mg/mL uridine (Sigma-Aldrich, U3003). Cells were plated on 0.1mg/mL poly-D-lysine (Sigma-Aldrich, P7405)-coated 12mm glass coverslips in a 24-well plate at a density of 150-200 neurons per coverslip, left to adhere at 37°C and 5% CO2 for 3 hours, then flooded with prewarmed media and maintained at 37°C and 5% CO2. Half media changes were performed every second day.

### Calcium imaging of human sensory neurons

Validation of potentially pain-promoting ligands upregulated in GBS was performed by calcium imaging of primary human DRG-derived sensory neurons. Coverslips were removed from media, washed briefly in bath solution (145 mM NaCl, 3 mM KCl, 2.5 mM CaCl_2_, 1.2 mM MgCl_2_, 10 mM HEPES, 7 mM glucose, pH 7.4±0.1, osmolarity 320±3), and then incubated with 3 µg/mL Fura-2 calcium indicator (Thermo Fisher Scientific, F1221) and 2% BSA in bath solution for 45 minutes at RT and maintained in the dark.

Fura-2 underwent de-esterification by incubating in standard bath solution for 30 minutes at RT. During this step, coverslips were additionally treated with recombinant human CCL4/MIP-1β (100ng/mL, PeproTech, 300-09) or its vehicle (0.1% BSA in PBS) for 30 minutes. For imaging, coverslips were placed on an inverted fluorescent microscope (Nikon Eclipse Ti2). Images were captured alternating between 340 nm and 380 nm of excitation every 0.5 seconds. Cells were perfused with bath solution for 90 seconds, then stimulated with 20 nM capsaicin (Sigma-Aldrich, M2028) for 2 minutes.

Capsaicin is an agonist of TRPV1 which is highly expressed on human nociceptors and at this concentration is appropriate for testing priming effects of other stimuli (99). Cells were then washed with bath solution for 2 minutes and then treated with 50 mM KCl for 1 minute to confirm neuronal viability. The 340/380nm signal ratio was calculated for each neuron and corrected for background signal using NIS elements software (Nikon, ver. 6.10.01). Neurons were considered viable and included in analysis if they displayed a ≥25% increase in the 340/380nm ratio following KCl stimulation and classified as a capsaicin responder if they further displayed a ≥25% increase following capsaicin stimulation. Statistical analysis was performed in *R* using Welch’s unequal variance *t*-tests as implemented in the rstatix package. Differences in responder rates between conditions were assessed using Fisher’s exact test.

### Statistics

All analysis was conducted in *R* (ver. 4.4.1) using RStudio, and all figures were produced in *R* except for iPathwayGuide outputs which were exported from the software. *P*-values were corrected for multiplicity of comparisons as stated. The threshold for statistical significance was set to α = 0.05.

### Study approval

All work was carried out in accordance with relevant laws, institutional guidelines, and the Declaration of Helsinki. The Baylor College of Medicine and University of Alabama at Birmingham Institutional Review Boards approved this study (protocol numbers H-22354 and F130930003 respectively). Written informed consent was obtained from all participants prior to phlebotomy, including permission to utilize samples for future genetics research.

## Data availability

The raw and processed sequencing data is available in NCBI’s Gene Expression Omnibus (101) and are accessible through GEO Series accession numbers GSE304871 (bulk RNA-seq) and GSE304872 (single-cell RNA-seq). Values for all data points in graphs are reported in the Supporting Data Values file.

## Author Contributions

JAO: Investigation, Software, Formal Analysis, Visualization, Writing – original draft; JBL: Investigation, Writing – original draft; IS: Investigation, Software, Writing – original draft; AAT: Investigation; NI: Software; SS: Investigation, Writing – review & editing; HM: Software; KES: Resources; TJP: Conceptualization, Methodology, Resources, Supervision, Funding acquisition, Writing – review & editing; EEU: Conceptualization, Methodology, Resources, Supervision, Funding acquisition, Writing – review & editing.

## Acknowledgements

The authors are grateful to the organ donors and their families for their gift, members of the Southwest Transplant Alliance for supporting the tissue recovery work, and members of the Price laboratory. The authors also acknowledge Dr. Yeunhee Kim and the UT Dallas Genome Core Research Facility for assistance with library preparation and RNA sequencing.

