## Supplementary material for "Nerve-targeting and pain-promoting transcriptomic signatures in early Guillain-Barré syndrome": Supllemental materials

**Supplemental Table 1:** Additional patient clinical information.

**Supplemental Table 2:** Bulk RNA-sequencing differential expression results.

**Supplemental Table 3:** Single-cell RNA-sequencing differential expression results.

**Supplemental Table 4:** Cell sorting antibody panel.

**Supplemental Table 5:** Bulk RNA sequencing quality control.

**Supplemental Table 6:** Number of cells included for single-cell RNA sequencing analysis.

**Supplemental Table 7:** Calcium imaging organ donor information.

**Supplemental Figure 1:** Single-cell feature dimensionality reduction.

**Supplemental Figure 2:** Single-cell feature dot plots.

**Supplemental Figure 3:** Classical and intermediate monocyte ligand-receptor analysis.

**Supplemental Figure 4:** FACS gating strategy.

**Supplemental Figure 5:** Single-cell quality control processing.

### Supplemental Table Captions

**Supplemental Table 1.** Additional clinical data for GBS and control subjects in the study as supplemental to main Table 1. HC4 was not included in any sequencing results.

**Supplemental Table 2.** Bulk RNA sequencing DESeq2 results for CD4+, CD8+, and CD11b+ sorted leukocyte subpopulations between GBS- and control-derived leukocytes.

**Supplemental Table 3.** Single-cell RNA sequencing DESeq2 results for GBS patient and control subject-derived leukocytes. The 22 sheets list the differential expression analysis for each gene across all clusters.

**Supplemental Table 4.** Reagent information for the fluorescence-activated cell sorting (FACS) preceding bulk RNA sequencing.

**Supplemental Table 5.** Quality control data corresponding to RNA extractions from CD4+, CD8+, and CD11b+ sorted cells from GBS and control blood preceding bulk RNA sequencing. RNA concentration, 260/280 and 260/230 ratios as determined by NanoDrop, RNA quality number, and gDNA contamination determined whether RNA samples were passed on for library preparation. Input cell counts are also provided.

**Supplemental Table 6.** Number of cells included for single-cell RNA sequencing analysis per sample.

**Supplemental Table 7.** Organ donor information for dorsal root ganglion sensory neuron calcium imaging experiments.

### Supplemental Figures

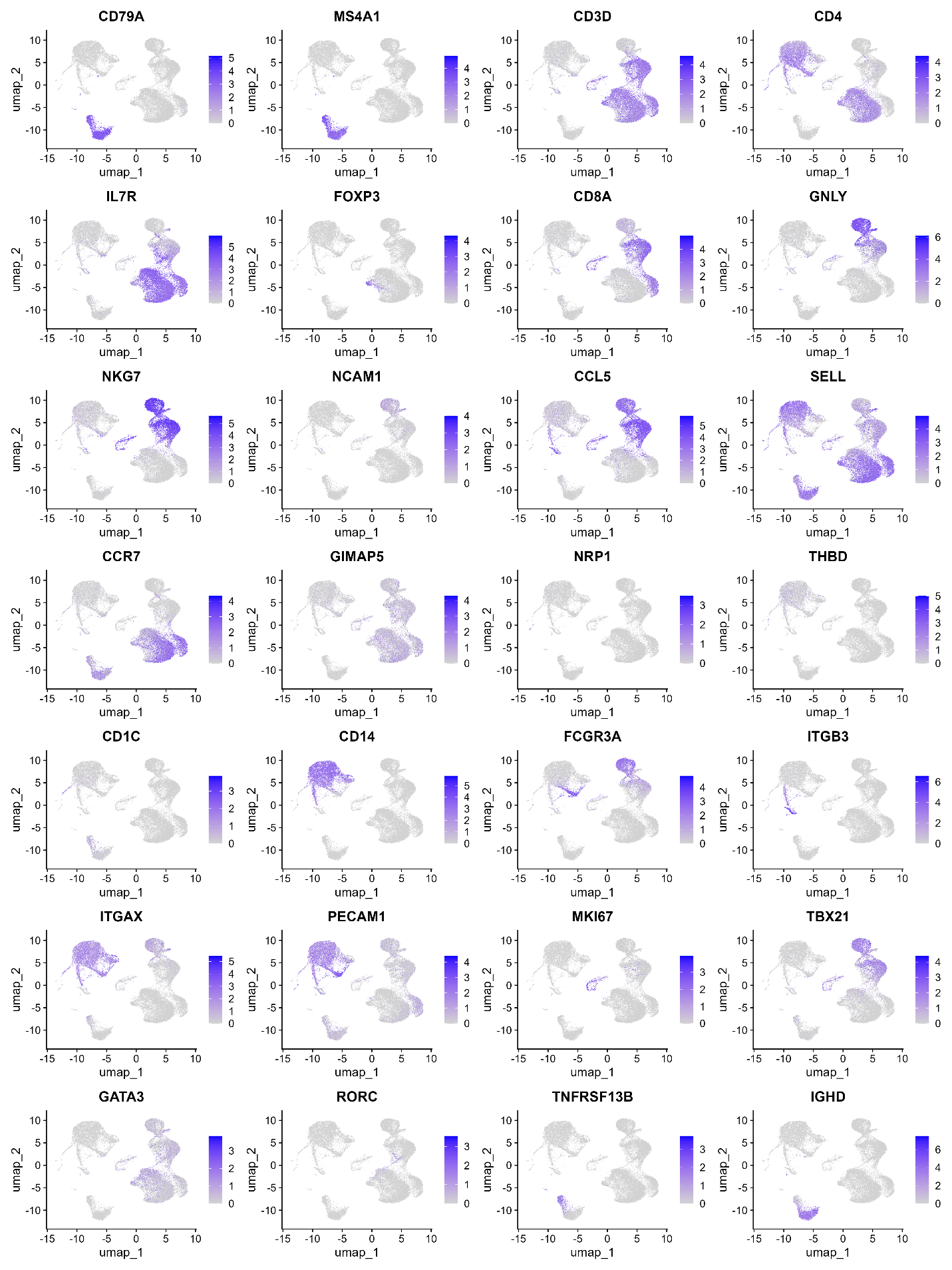

**Supplemental Figure 1.** UMAP of key cell type-specifying genes.

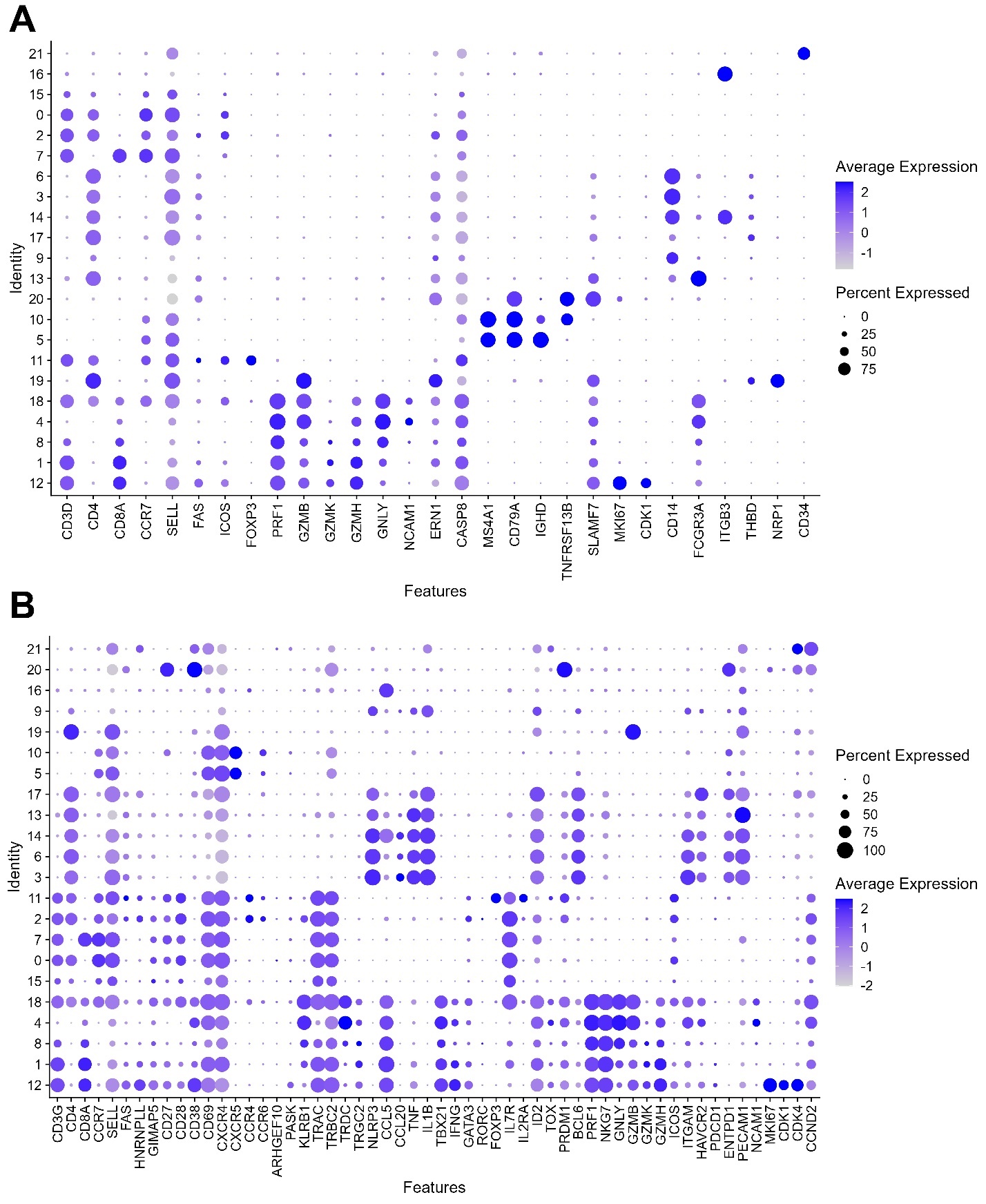

**Supplemental Figure 2.** Dot plots of key cell type-specifying genes used for cluster annotation. **(A)** Expression within each cluster of genes corresponding to major cell types. **(B)** Expression within each cluster of genes corresponding to T cell differentiation, activation, and function.

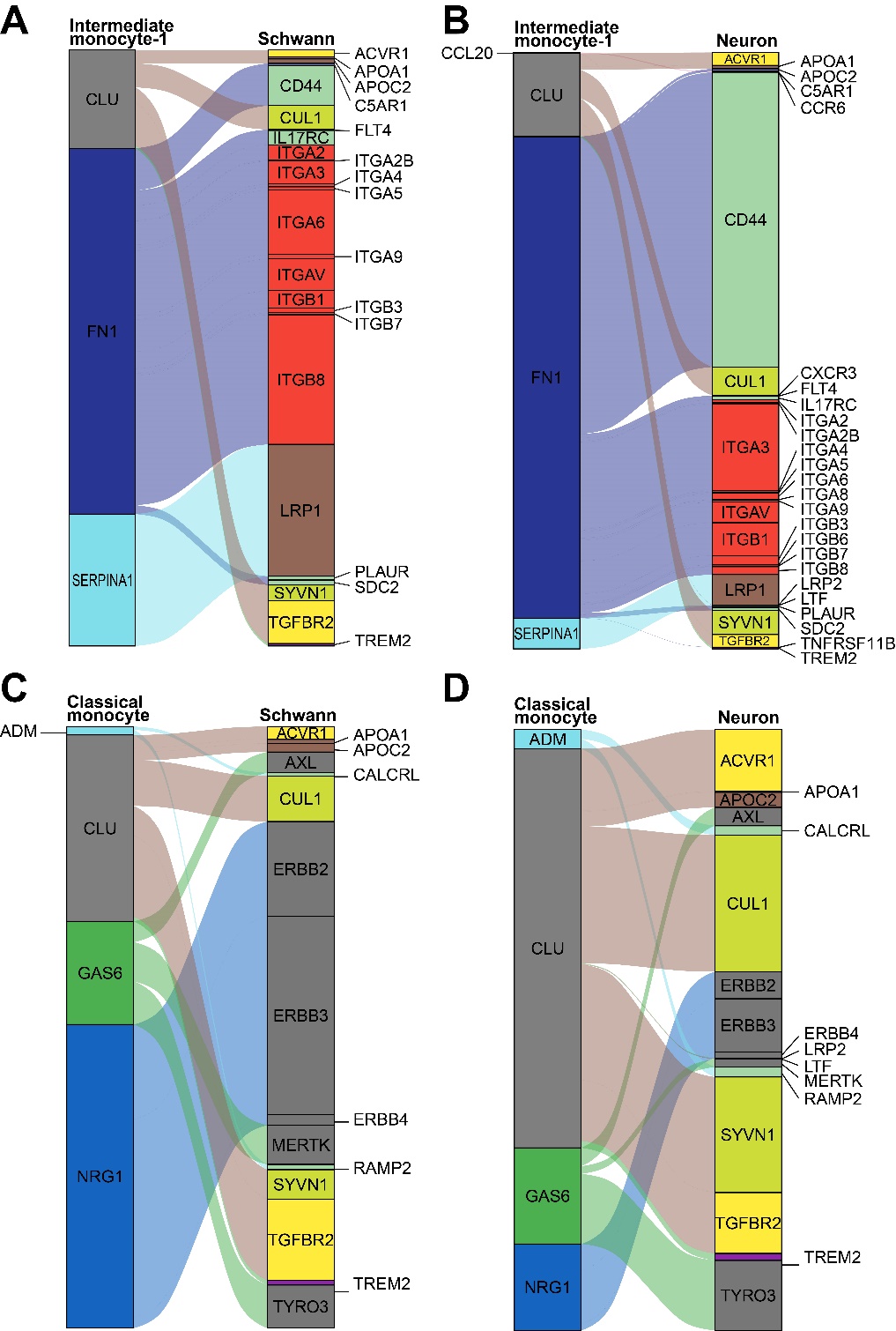

**Supplemental Figure 3.** Ligand-receptor pairs forming predicted interactions between upregulated ligand genes in GBS patient intermediate and classical monocytes toward receptors expressed in human dorsal root ganglion Schwann cells and sensory neurons. **(A)** Interactions between intermediate monocyte-1 (cluster 6) ligands upregulated in GBS and myelinating Schwann cells and **(B) sensory** neurons. **(C)** GBS patient upregulated genes in classical monocyte (cluster 3) cells and their predicted interactions with receptors on myelinating Schwann cells and **(D)** sensory neurons. Boxes for each gene are weighted by expression. Alluvial flows are weighted by the expression of the relevant receptor.

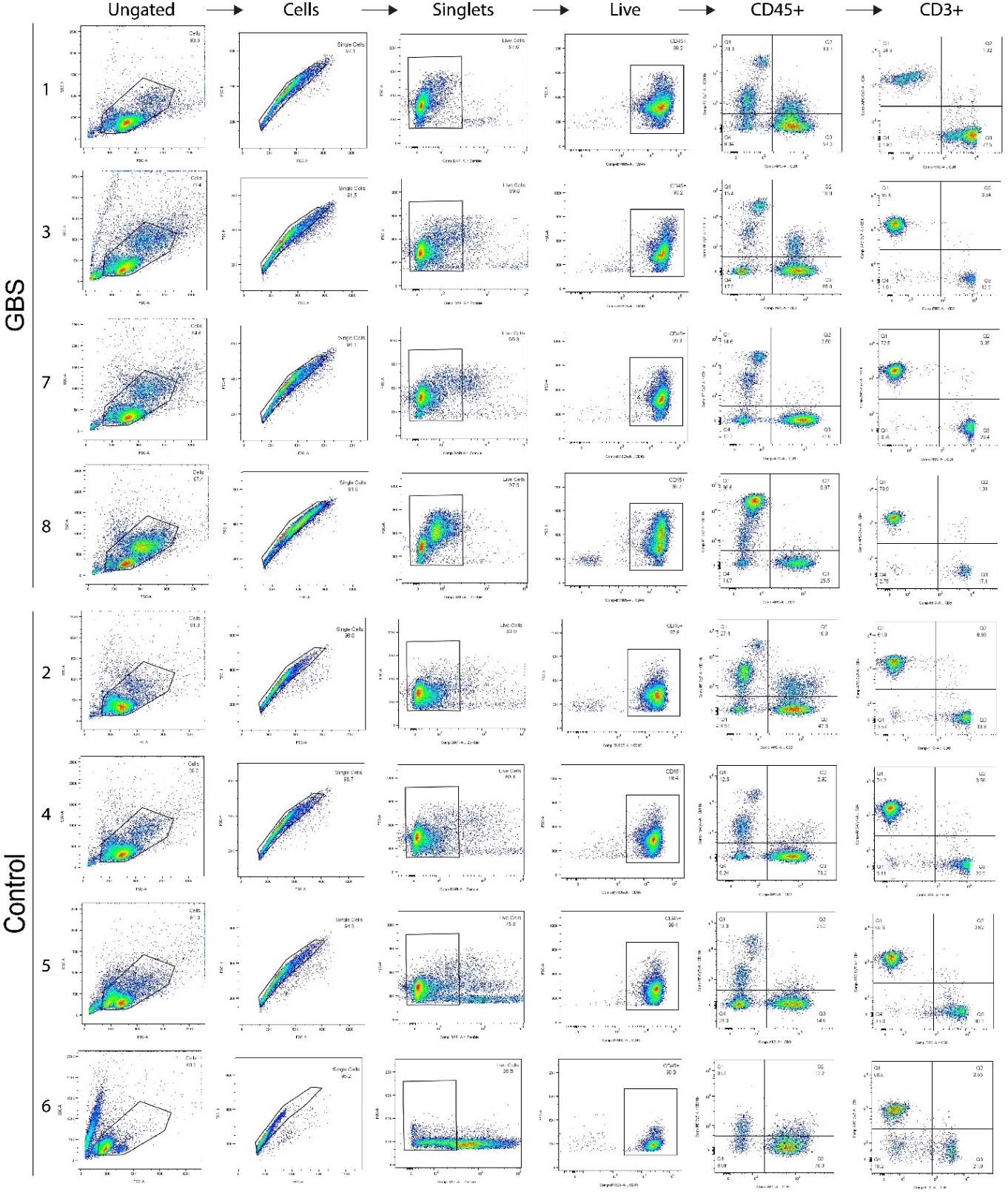

**Supplemental Figure 4.** Gating strategy for fluorescence-activated cell sorting of GBS and healthy control peripheral blood mononuclear cells by sample number. These gates were used to generate the CD4+, CD8+, and CD11b+ cell populations for onward RNA extraction and bulk RNA sequencing, while also providing quality control data for single-cell sequencing. Sample #6 was excluded due to a high proportion of dead cells.

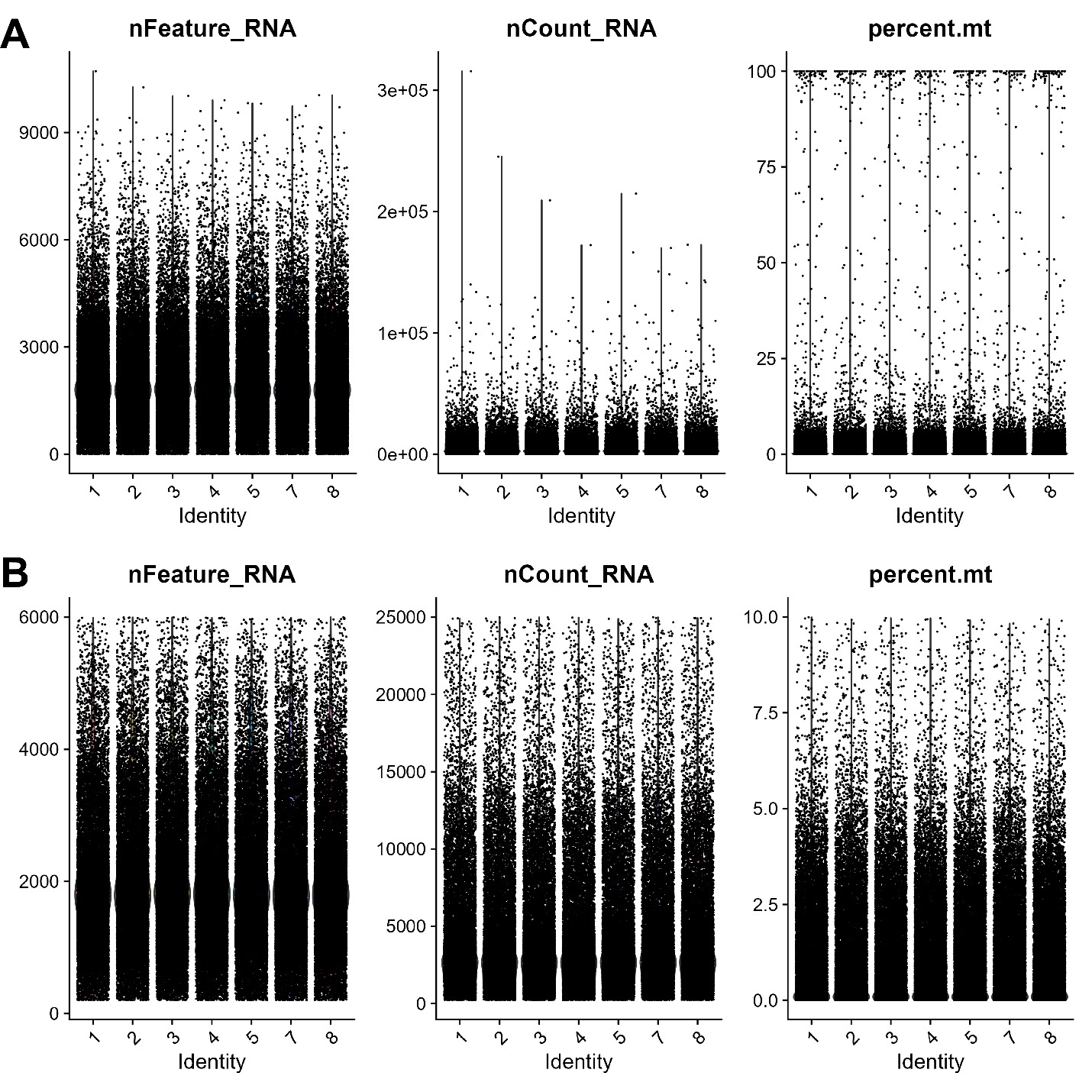

**Supplemental Figure 5.** Quality control measures for single-cell RNA sequencing of control and GBS patient-derived PBMCs. **(A)** The number of features, total counts, and percentage of mitochondrial genes in each cell before filtering. **(B)** The same parameters after filtering out cells with more than 6000 features, 25000 counts, or 10% mitochondrial genes.
